# Clonal dynamics in early human embryogenesis inferred from somatic mutation

**DOI:** 10.1101/2020.11.23.395244

**Authors:** Seongyeol Park, Nanda Maya Mali, Ryul Kim, Jeong-Woo Choi, Junehawk Lee, Joonoh Lim, Jung Min Park, Jung Woo Park, Donghyun Kim, Taewoo Kim, Kijong Yi, June Hyug Choi, Seong Gyu Kwon, Joo Hee Hong, Jeonghwan Youk, Yohan An, Su Yeon Kim, Moonkyu Kim, Dong Sun Kim, Ji Young Park, Ji Won Oh, Young Seok Ju

## Abstract

The trillions of cells that constitute the human body are developed from a fertilized egg through embryogenesis. However, cellular dynamics and developmental outcomes of embryonic cells in humans remain to be largely unknown due to the technical and ethical challenges. Here, we explored whole-genomes of 334 single-cell expanded clones and targeted deep-sequences of 379 bulk tissues obtained from various anatomical locations from seven individuals. Using the discovered 1,688,652 somatic mutations as an intrinsic barcode, we reconstructed cellular phylogenetic trees that provide novel insights into early human embryogenesis. Our findings suggest (1) endogenous mutational rate that is higher in the first cell division of life but decreases to ~1 per cell per cell division later in life, (2) universal unequal contribution of early cells into embryo proper resulting from early cellular bottlenecks that stochastically separate epiblasts from embryonic cells (3) uneven differential outcomes of early cells into three germ layers, left-right and cranio-caudal tissues, (4) emergence of a few ancestral cells that will contribute to the substantial fraction of adult blood cells, and (5) presence of mitochondrial DNA heteroplasmy in the fertilized egg. Our approach additionally provides insights into the age-related mutational processes including UV-mediated mutagenesis and loss of chromosome X or Y in normal somatic cells. Taken together, this study scrutinized somatic mosaicism, clonal architecture, and cellular dynamics in human embryogenesis at an unprecedented level and provides a foundation for future studies to complete cellular phylogenies in human embryogenesis.

## Introduction

An adult human body consists of trillions of cells of more than 200 types^1^. These cells can be traced back to the fertilized egg, which continues to divide and eventually organizes into an individual body. Most of our knowledge about human development originates from rare direct observation of human fetal specimens, or extrapolation from model organisms despite the interspecies divergence^2–6^. Despite extensive studies^7,8^, our understanding of the fundamental aspects of early human embryogenesis remains limited.

Genomic mutations spontaneously accumulate in somatic cells throughout life^9^, presumably to a substantial rate from the first cell division^10^ by intrinsic mechanisms or exposure to external mutagens^11^. As the vast majority of mutations arising in an ancestral cell are inherited by its descendant cells, these mutations can be used as cellular barcodes to reconstruct developmental phylogenies of somatic cells^12^, as used in tracking the clonal evolution in tumor tissues^13,14^, normal blood^14^ and brain development^15,16^. To reconstruct early lineages of somatic cells in an individual, two experimental conditions are necessary: (1) sensitive and precise discovery of mutations acquired during the early embryogenesis, and (2) comprehensive tracing of the mutations (or the mutant cells) in various human tissues. To this end, our study applied the ‘capture-recapture’ strategy with early somatic mutations (**Fig. 1a**) ^15^. Together with a concurrent study using a different but complementary approach, we explored the clonal dynamics in the early human embryogenesis across individuals.

**Fig 1.**
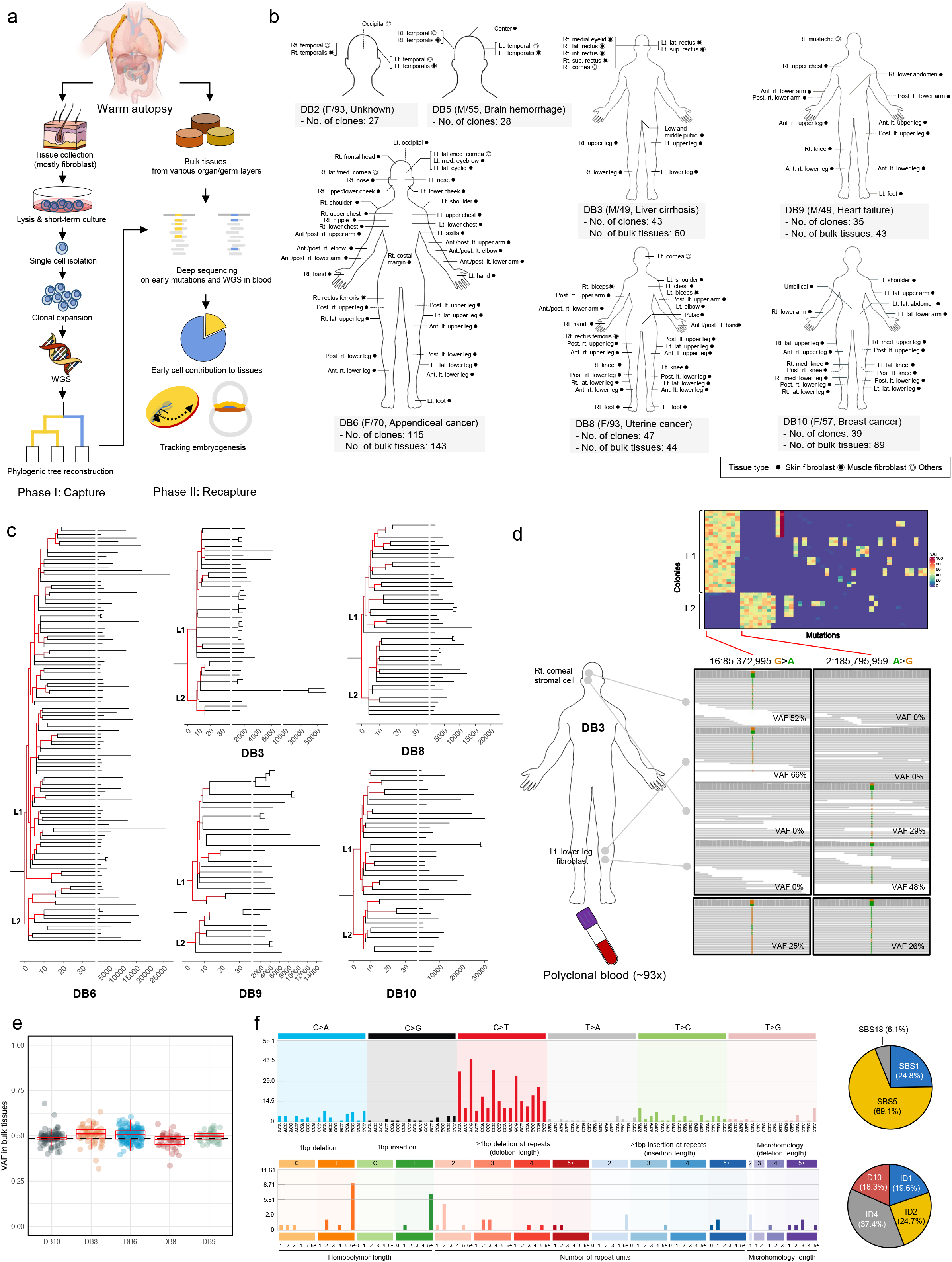
Reconstruction of the early embryogenic cellular phylogenies using somatic mutations. **a,** experimental design with two phases: a capture phase, in which adult viable single cells were collected and expanded *in vitro* and whole-genome sequenced for detecting genome-wide somatic mutations, and a recapture phase, in which bulk tissues were deep-sequenced for estimating the cellular frequency of the early embryonic mutations discovered in the capture phase. **b,** summary of the seven warm autopsies, in which clones and bulk tissues were collected. **c,** the phylogenetic trees of the clones showing early embryonic cell divisions. Horizontal axes are molecular time by mutation numbers. The branch lengths are proportional to the number of somatic mutations. Branches with defined early mutations were colored in red. Two initial branches are marked with L1 (major contribution) and L2 (minor contribution) according to their relative contribution in clones. **d,** examples of embryonic mutations found in the whole-genome sequencing of the clones and polyclonal blood cells. A heatmap in the upper panel shows the variant allele fractions (VAFs) of the early embryonic mutations in the clones of an individual (DB3). IGV screenshots for the two early embryonic mutations in WGSs of four clones and polyclonal blood are also shown. **e,** the aggregated VAFs for the first two mutation subsets in the bulk tissues (dot) showing a very close distribution from 0.5. **f,** the signatures of the early embryonic substitutions (n=488) and indels (n=49) which are delineated by COSMIC signatures^56^.

### Capture-recapture of early embryonic mutations

First, in the ‘capture’ phase, we isolated single cells from various anatomical locations across entire human bodies and explored somatic mutations harbored in these cells (**Fig. 1b; Supplementary Discussion 1**). Although single-cell genome sequencing is conceptually straightforward, it is often associated with a myriad of challenges, including high error rates and allele drop out during whole-genome amplification, which significantly hinder sensitive and specific discovery of somatic mutations^17^. To overcome the issues, we clonally expanded single cells *in vitro* to ~1,000-10,000 cells, which provide a sufficient amount of genomic DNA (~6-60 ng) for conducting amplification-free whole-genome sequencing (WGS). Viable cells, mostly skin and muscle fibroblasts, were obtained from seven individuals at autopsy (**Supplementary Table 1**). Overall, we established 374 colonies from the seven cases followed by WGS to 25x on average. After excluding colonies with low-coverage WGS data or established from multiple founder cells (**Extended Data Fig. 1a**), we obtained high quality WGS data from 334 single-cell colonies (hereafter referred to as clones) from the seven donors (279 clones from whole-body locations from five donors (DBs 3,6,8,9 and 10) and 55 clones from limited sites (head regions) from two further donors (DBs 2 and 5)) (**Supplementary Table 2**). Through extensive downstream bioinformatic analyses, all types of somatic mutations, including base substitutions, indels, copy number changes, and structural variations, were comprehensively explored. By comparison of somatic mutations across multiple clones, we systematically identified mutations acquired in the common ancestral cells of the clones, mostly early in embryogenesis.

Second, in the ‘recapture’ phase, we dissected small bulk tissues (n=379) from various organs and anatomical locations encompassing three germ layers in five individuals (DBs 3, 6, 8, 9, and 10) (**Fig. 1a; Supplementary Table 3**). The recapture was not possible from the other two individuals (DBs 2 and 5) due to the limited samples available. Where possible, skin bulk tissues were further separated into epidermis and dermis to differentially explore ectodermal and mesodermal cell pools (**Methods**). Deep sequencing (mean ~911x) was performed on the bulk tissues targeting early mutations discovered in the ‘capture’ phase to trace the contribution of each variant to the adult tissues. In addition, we also deeply sequenced whole-genome of polyclonal blood samples from each individual (mean depth of ~100x) to track early cells in the hematopoietic tissue.

### Somatic mutations and early phylogenies

From WGS of the 334 clones, we identified overall 1,647,684 somatic single nucleotide variants (SNVs, or base substitutions) and 40,968 small insertions and deletions (indels). Four levels of evidence indicated that the vast majority of our calls were true somatic mutations accumulated *in vivo*, rather than culture-mediated artifical mutations. First, variant-allele frequencies (VAFs) were distributed around 50% in WGS (**Extended Data Fig. 1b, c**), suggesting that these were harbored in all cells in the clones. Second, mutational spectrums were delineated by biological mutational signatures rather than signatures for sequencing artifacts (mutational signatures will be discussed later) ^18^. Third, we experimentally confirmed the rate of culture-associated mutations in three pairs of serial single-cell expansions (**Extended Data Figs. 1d - f**). The results showed that, on average, 7.6% (5.0% - 9.3%) of the mutations in a clone could be culture-associated. Lastly, in clones derived from male donors, only 3.3% of mutations in the X chromosome exhibited heterozygous-like patterns (VAF < 0.7), which could be subclonal and potentially culture-associated mutations.

For reconstruction of early phylogenies, we used WGS data from 279 colonies derived from whole-body locations from five donors (DBs 3, 6, 8, 9, and 10). Overall, a total of 1,532,801 somatic SNVs and 35,258 indels were identified (**Supplementary Table 2**). Of these, 117,342 SNVs and 2,405 indels were shared by two or more clones, suggesting that these mutations were present in the most recent common ancestral cells of the clones thus enabling reconstruction of the cellular phylogenies (**Extended Data Fig. 2**). Intriguingly, a small set of mutations was sufficient for each individual to completely partition all the clones into two groups (referred to as L1 and L2 lineages) (**Fig. 1c**). For example, chr16:85,372,995 G>A substitution and 7 other mutations were present in 30 of the 43 clones in DB3 (L1), while chr2:185,795,959 A>G substitution and another set of 7 mutations were present in all the remaining clones (L2) (**Fig. 1d**). Two sets of mutations defining the L1 and L2 lineages were mutually exclusive in all five individuals: no clones possessed both or neither of the mutation sets (**Extended Data Fig. 3a**). The two sets of mutations were found in all the bulk polyclonal tissues from blood and other solid tissues with aggregated L1 and L2 VAFs to be ~50% (**Fig. 1e**). It suggests that these mutations were cellular markers for the two earliest ancestral cells of all somatic cells in DB3, potentially, but not necessarily, two cells at the two-cell stage embryo. After the first cell branching, clones were further partitioned by additional mutations shared by a smaller subgroup of cells (**Fig. 1c; Supplementary Table 4**). Except for a few outlier clone pairs sharing several hundred to thousands of mutations, which likely branched from a common ancestral stem cell in an aged adult tissue, most clone pairs diverged from each other before 35 mutations, which defined the latest embryonic period possible to explore in this study (**Fig. 1c**).

Our phylogenetic trees indicated that anatomically adjacent cells in the adult tissues are not always developmentally closer. For example, a fibroblast clone established from the left lower leg shared the same L2 mutations with the right corneal stroma, while another lower leg clone derived from a cell in its vicinity was a progeny from the L1 lineage (**Fig. 1d**).

From the five developmental phylogenetic trees, 488 SNVs and 49 indels were acquired earlier than 35 mutations, our latest embryonic period (**Fig. 1c**); therefore, they are defined as early embryonic mutations in this study. As expected, the number of detected early mutations per individual increases with the number of clones sequenced (**Extended Data Fig. 3b**). Our data directly show that indels can accumulate from early embryogenesis, with an ~10% rate of base substitution, as typically observed in aged cancer cells (**Extended Data Fig. 3c**) ^19^. The mutational spectrums suggest that previously defined mutational signatures of endogenous processes from single-base substitutions (SBSs) and indels (IDs) ^18^, such as SBS1 (5-methylcytosine deamination) and SBS5 (unknown etiology) and ID4 (unknown etiology) as well as ID1, ID2 (both from slippage during DNA replication), are dominantly responsible for early mutations (**Fig. 1f**)(see also COSMIC database (https://cancer.sanger.ac.uk/cosmic/signatures) for the latest mutational signatures). Compared to adult-stage mutations, we found a higher fraction of SBS1 in early mutations relative to SBS5 (odds ratio = 3.7, p < 0.001; **Extended Data Fig. 3d**), which implies a higher cell division rate in early embryogenesis. Mutational signatures for indels were also different between early and late stages (**Extended Data Fig. 3d**).

### Features of early developmental phylogenies

We observed two common characteristics elucidating early clonal dynamics from the five developmental phylogenetic trees with whole-body clones. First, unlike absolute bifurcations (dichotomies) in the first branchings, multifurcations (polytomies) were often observed from the second branchings (**Fig. 1c, Extended Data Fig. 4a**). The transition from early dichotomies to late polytomies is consistent with three other autopsies in the companion paper. Collectively, our trees imply that the mutation rate in the first cell branching in life is higher than that of the subsequent branchings. It also points to that subsequent cell branchings are not always ‘informative’ in revealing physical cell divisions due to the stochastic absence of intrinsic mutation (**Extended Data Fig. 4b, c**).

Second, as observed previously^10^, the two earliest ancestral cells (and other branchings afterward) showed an unequal contribution in the phylogenetic trees (**Fig. 2a**). For instance, 112 lineages in DB6 split at a ratio of 6.5:1 (86.6% vs 13.4%), substantially skewed from the 1:1 expectation (p < 0.001). The asymmetry was undoubtedly not an illusion caused by sampling bias in our single-cell clonalization, because VAFs of L1 mutations are constantly higher than those of L2 mutations in the targeted deep sequencing of bulk tissues (**Fig. 2a**). In agreement with DB6, unequal contributions were consistently observed in the first branchings of other phylogenies, but the asymmetry levels were variable among individuals in the range of 1.4:1 to 6.5:1 ratio. The unequal contribution between two daughter cells extends to the second and later branchings in most early embryonic cells bifurcated in the trees (**Fig. 2a, right**), which were concordant with VAFs from targeted deep sequencing of various bulk tissues (**Fig. 2b**). These suggest that our clones established from whole-bodies well reflect the true clonal composition of the somatic cells.

**Fig 2.**
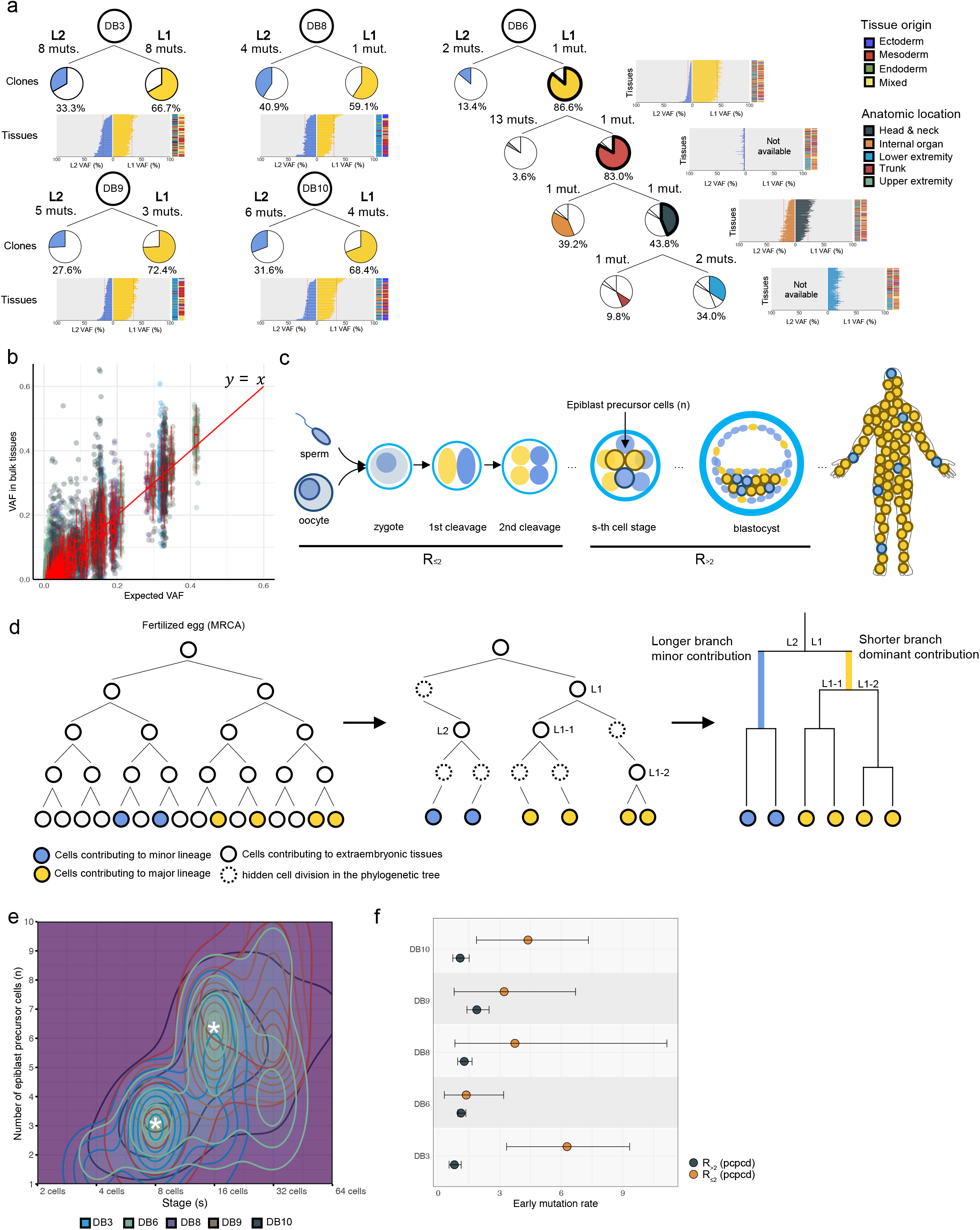
Universal unequal contribution of early embryonic cells to adult tissues with variable asymmetry across individuals. **a,** unequal contribution of the two earliest branches (L1 and L2) consistently found in phylogenies and bulk tissues. Asymmetry between bifurcating downstream branches were further tracked in DB6. Pie graphs represent the contribution of each branch to lineages (clones) in phylogenetic trees. Horizontal bar graphs show the VAFs of the lineage-specific mutations in bulk tissues. Expected VAFs from the phylogenies are shown by red dashed lines. **b,** VAFs of the early embryonic mutations in bulk tissues (*y* axis) are collectively concordant with the expected from the phylogenetic trees (*x* axis). **c,** a developmental model with the lineage asymmetry structured in the early cell pool, which can explain the unequal contribution conserved in most adult tissues of an individual body. This model assumes the number of cells (*n*) are selected for the epiblasts formation at a certain cell stage (*s*). We presumed two different mutation rates (R_<2_ and R>2) in the early embryogenesis. **d,** a cellular genealogy scenario demonstrating a developmental cellular bottleneck that can shape the asymmetric features observed in the phylogenetic trees. **e,** a contour plot showing the best-fitting simulation considering a developmental bottleneck. The horizontal and vertical axes represent the absolute embryonic cell stage and the number of contributing cells to the pool of epiblasts. White asterisks indicate the values of the maximum likelihood. **f,** estimates of early embryonic mutation rates per cell per cell division (pcpcd), considering a developmental bottleneck and potential missing nodes. R_<2_, mutation rate until the 4-cell stage, R_≥2_, mutation rate from the 8-cell stage.

Dominant and minor clonal relationships universally observed in most whole-body tissues of an individual body suggest that the asymmetry was structured in the early founder cells of human somatic cells, the inner cell mass (ICM) or epiblasts, which eventually contribute to all somatic cells (**Fig. 2c**) ^20^. Simultaneously, variable levels of unequal contribution across the individuals suggest that some extent of stochasticity is involved in the unequal cellular segregation process, unlike *C. elegans* with a highly deterministic embryogenesis (**Fig. 2a, Extended Data Fig. 4d**) ^21,22^.

To understand the biological basis underlying the unequal contribution in the early cellular phylogenies, we employed known biological contexts of early embryogenesis. Briefly, in the blastocyst-stage human embryo, a few minority cells are segregated (or selected) through biological bottlenecks to the ICM and then to epiblasts. In this circumstance, some early cells may not contribute to the embryo proper but constitute extraembryonic tissues exclusively; therefore, they are missing from our ‘mutational capture’ phase because these cells cannot contribute to somatic clones. As illustrated in **Fig. 2d**, the ‘missing embryonic cells’ have two impacts: (1) progeny cells in the lineage harboring many missing cells will contribute less in the embryo proper and will be minor in the phylogenetic tree and the adult tissue, and (2) two or more consecutive cell divisions will coalesce into one cell branch in the phylogenies, thus the cell doubling will more likely engage a longer molecular time (more cell divisions and thus more mutations). Indeed, early bifurcating nodes in the five trees showed the trend; the branches with a relatively smaller cellular contribution have the same or longer lengths (13 out of 19 bifurcating nodes; 68.4%; **Fig. 2a, Extended Data Fig. 4a**).

For more quantitative investigation, we simulated a cell genealogy scenario in which a few cells are stochastically set to epiblast precursor cells through a cellular bottleneck in a fully bifurcating tree at a certain cell stage (**Fig. 2c, Methods**). With our phylogenies and number of early somatic mutations, our simulation studies together with likelihood approach inferred that three cells in the 8 cell-stage embryos, or six cells in the 16 cell-stage embryos, are main ancestral cells contributing to the epiblasts, suggesting that symmetry of early cells could be broken at the stage (**Fig. 2e**). Our conclusion agrees with previous mouse studies^23^ and the divergence of the trophoblast-derived tissue from blood antecedents at the 4 to 16 cell-stage human embryo^24^.

By accommodating missing cell divisions in the early phylogenetic trees, our simulation efforts determined early mutation rate per absolute cell division. With an assumption of two different early mutation rates changing at 4-cell stage embryos as we observed in the phylogenetic trees, the maximum likelihood estimations of the early mutation rates were ~3.8 (range 1.4 to 6.3) mutations per cell per cell division (pcpcd) for the earliest cell divisions (R_<2_) and ~1.2 mutations pcpcd (range 0.8 to 1.9) from 8-cell stages and after (R_≥2_) (**Fig. 2f**). Of note, zygotic genome activation (ZGA) occurs near the transition period^25,26^, and we speculate that the higher mutation rate in the earlier period may be, in part, due to the lack of mature DNA repair processes in the pre-ZGA embryos. Of note, the marginal mutation rate for the later early cell divisions (near 1 pcpcd) is in agreement with the relatively frequent multifurcation from the second branching.

### Tracing the developmental outcomes of early embryonic cells

By exploring the VAFs of the early embryonic mutations in deep sequencing (recapture), we traced the contribution of early cells in various adult tissues. Intriguingly, from molecular time of ~9-15 mutations, the VAFs of early mutations were consistently lower in ectodermal tissues (e.g., epidermis) than those from the mesodermal (e.g., dermis and muscle) or endodermal tissues (e.g., liver and lung) in all available cases (**Fig. 3a**). It implies that a few lineages specifically contributing to ectodermal tissues diverged from lineages that predominantly contributing, but not limited, to meso- and endodermal tissues. By marking mutations with different VAFs on the phylogenetic tree of DB6, we estimated that the ectoderm-specific cells diverged around the 16-cell stage in this individual (asterisk in **Fig. 3b**). Undifferentiated VAFs between the mesodermal and endodermal tissues suggest that these two germ layers were not committed until a later time period (**Fig. 3a**).

**Fig 3.**
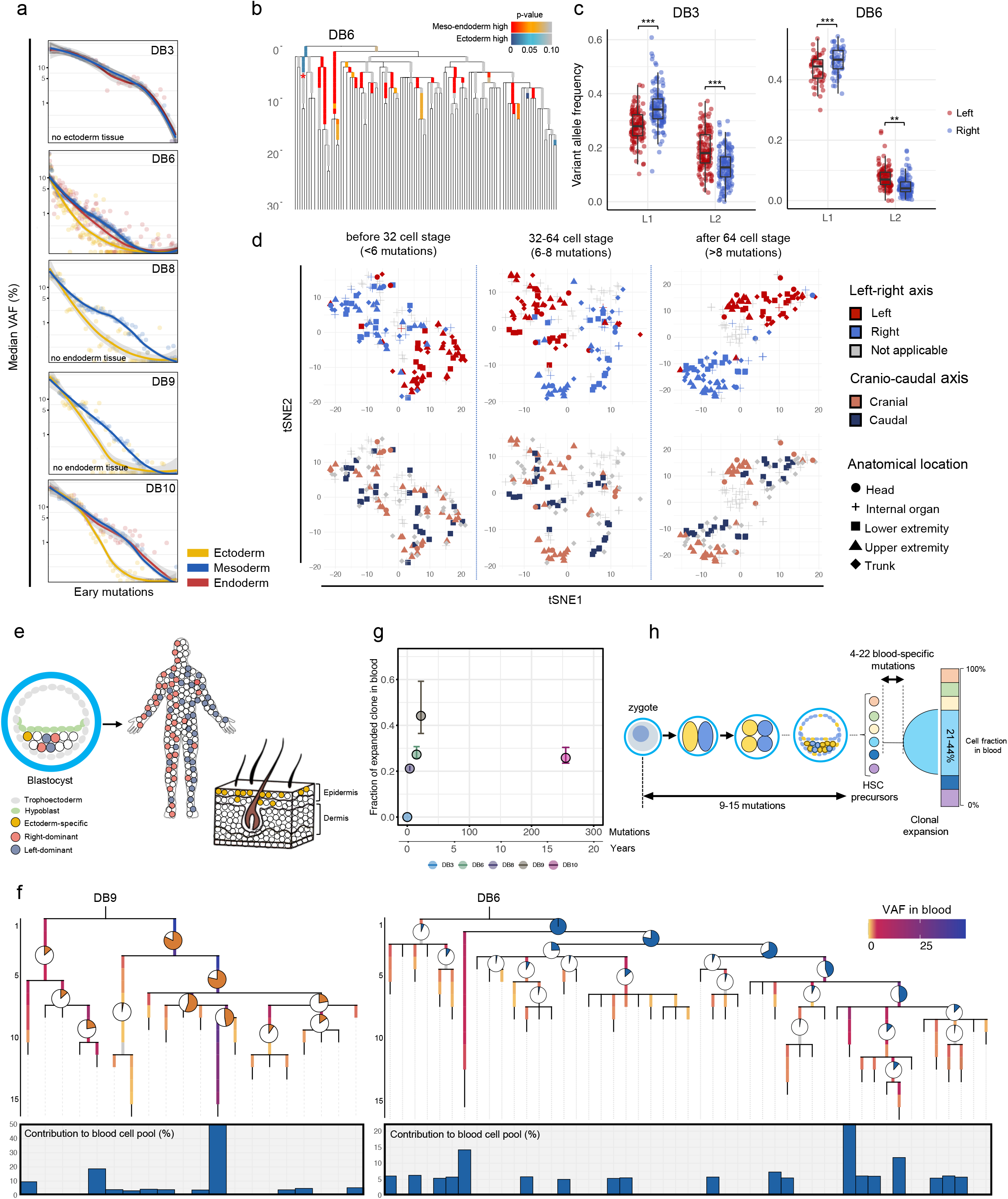
Timing of early cell fate determination into different tissues. **a,** Median variant allele frequencies (VAFs) of the early embryonic mutations in deep sequencing of the bulk tissues according to the dominant germ layers. The horizontal axis shows early mutations sorted by the averaged VAFs in bulk tissues in a descending order, thus from earlier to later mutations. The VAFs in the ectodermal, mesodermal, and endodermal tissues are colored in orange, blue, red, respectively. The lines are fitted curves by locally estimated scatterplot smoothing (LOESS) methods. **b,** the phylogenetic tree of DB6 showing asymmetric distribution between ectoderm and meso-endoderm. Mutations with lower VAF in ectoderm are colored by red, while ones with higher VAF in ectoderm are colored by blue. Comparisons were performed by Wilcox tests. A red asterisk indicates the branching out of the ectoderm-specific lineage. **c,** scatter plots illustrating the unequal contribution of the first two lineages to left- and right-side of body in DB3 (left) and DB6 (right). **d,** tSNE plots of DB6 bulk tissues using the VAFs of the embryonic mutations occurred in different molecular times (**Methods**). The separation between left and right tissues was apparent from the earliest molecular time (i.e. <6 mutations, top row) followed by the separation according to the cranio-caudal axis (bottom row). **e,** a schematic presentation demonstrating the unequal developmental outcomes of early cells in the epiblast. **f,** the emergence of an early embryonic clone dominantly contributing to the adult blood tissues from DB9 (left) and DB6 (right). Pie graphs (top, internal branches) and bar graphs (bottom, terminal branches) represent the averaged VAF of the mutations in the branch observed in the deep whole-genome sequencing of blood tissues. **g,** the numbers of blood-specific somatic mutations (horizontal axis) and their contributions to the total blood cells (vertical axis; vertical lines indicate interquartile ranges). Estimated age was calculated using 15 mutations/year in blood cells reported in the previous studies^15,27^. **h,** a schematic representation demonstrating the emergence of precursors for hematopoietic stem cells (HSCs) and their clonal expansions during embryogenesis.

In addition, we compared the VAFs of early mutations in left-vs. right-side tissues (left-right axis) and in upper-vs. lower-body tissues (cranio-caudal axis). The systematic differences across these axes were observed from the mutations acquired in the very early period. For example, we found a slight but significant asymmetry of the left-right axis from the first two lineages (L1 and L2) in DB3 and DB6 (**Fig. 3c**), suggesting that the earliest two ancestral cells of human bodies contributed unequally according to the axis. Despite slight individual variations in the amount and timing, all five cases exhibited asymmetries on the left-right and cranio-caudal axes from the embryonic periods we explored. Interestingly, the left-right asymmetry appeared earlier than the cranio-caudal asymmetry, i.e. molecular time of ~5 mutations in DB6 (which is equivalent to ~4 cell divisions) (**Fig. 3d and Extended Data Fig. 5a**). The early restriction of lateral cellular movement can be partially explained by the primitive streak separating the left and right of the epiblast layer in pre-gastrulation embryos (**Extended Data Fig. 5b**). Unequal developmental outcomes of early cells through the axis and in ectodermal tissues are conceptually summarized in **Fig. 3e**.

The contribution of early cells into hematopoietic tissues can be traced to a later molecular time (mutations occurred in the later cell divisions) using deep WGS (~100x) of polyclonal blood tissues. We found a few later lineages preferentially producing hematopoietic stem cells (HSCs) emerging at a certain time point. In DB9, for example, a lineage branched at a molecular time of 12.5 mutations contributed to ~50% of the adult blood tissue, which is 15-fold higher than the equal contribution from clones in the phylogenetic tree (**Fig. 3f)**. We consistently found these ‘blood-contributing’ cell lineages in all five individuals at a molecular time of ~9-15 mutations (equivalent to ~2nd - 6th cell divisions; **Extended Data Fig. 5c**). We can further determine the molecular time of the expansion of ‘blood-contributing single clone’ (which contributed 21-44% of all the blood cells) by detecting mutations in the blood WGS with substantial VAFs (**Fig. 3g, Extended Data Fig. 5d**). The numbers of blood-specific mutations were only 4 to 22, suggesting that the clones expanded in blood tissues after a few more cell divisions from early embryogenesis (except for DB10 with 255 mutations, estimated to be expanded ~17 years of age given ~15 mutations per year in HSCs^15,27^) (**Fig. 3g, 3h; Extended Data Fig. 5d**). Because of its relatively lower number of mutations, the early clones preferentially contributing to blood are distinguished from neoplastic clonal expansion of adult HSCs known as age-related clonal hematopoiesis^28^.

### Heteroplasmy of mitochondrial DNA

Multiple clones established from single cells in an individual enabled dissection of the mitochondrial DNA (mtDNA) heteroplasmy present in fertilized eggs. From the WGS of 279 clones from the five autopsies, we identified 485 somatic-like SNVs and 48 indels in the mtDNA, which are heterogeneously present among the clones of an individual. Most of them (84.4%) were found in a single clone at various levels of heteroplasmy, consistent with the expected more-or-less random acquisition of somatic mutations in each lineage. However, a fraction of mutations (18.0%; 96/533) were found recurrently in multiple intra-individual clones, suggesting that these are unlikely to be independent and recurrent mutations, but stemmed from a certain level of heteroplasmy present in their ancestral cell. For example, mtDNA:16,256 C>T substitution was found in 14 clones from DB10 (36.8% out of a total 38 clones), often to homoplasmic levels (~100% VAF) within the 11 clones (**Fig. 4a**). However, the remaining clones (63.2%; 24/38) in DB10 and clones established from other donors (n=241), did not show any evidence for the mtDNA mutation (**Extended Data Fig. 6a**). The variant was also observed in the WGS of polyclonal blood cells (with an aggregated heteroplasmy of 35.0%), reassuring that the mutation is biological. Simulation using the phylogenetic relationships of the mutant cells in the tree indicated that the ancestral mutant is the fertilized egg with an initial heteroplasmic level of ~29.8% (95% CI 25.0-34.0%; **Methods; Extended Data Fig. 6b-d**). We speculate that the mutation was mostly fixed (100% VAF), or lost (0% VAF), in the aged somatic cells by mitochondrial bottlenecks during a progressive mtDNA copy number reduction in the cleavage^29,30^, and/or by random-walk mtDNA segregations through thousands of cell divisions for a lifetime operative in each cellular lineage (**Fig. 4b**) ^31–33^. Albeit at lower heteroplasmic levels, we found mtDNA heteroplasmy in the first cell from the other donors (**Fig. 4c**), suggesting that heteroplasmic DNA mutations are not rare in oocytes despite the mtDNA purification mechanism in the maternal germline^34–36^.

**Fig 4.**
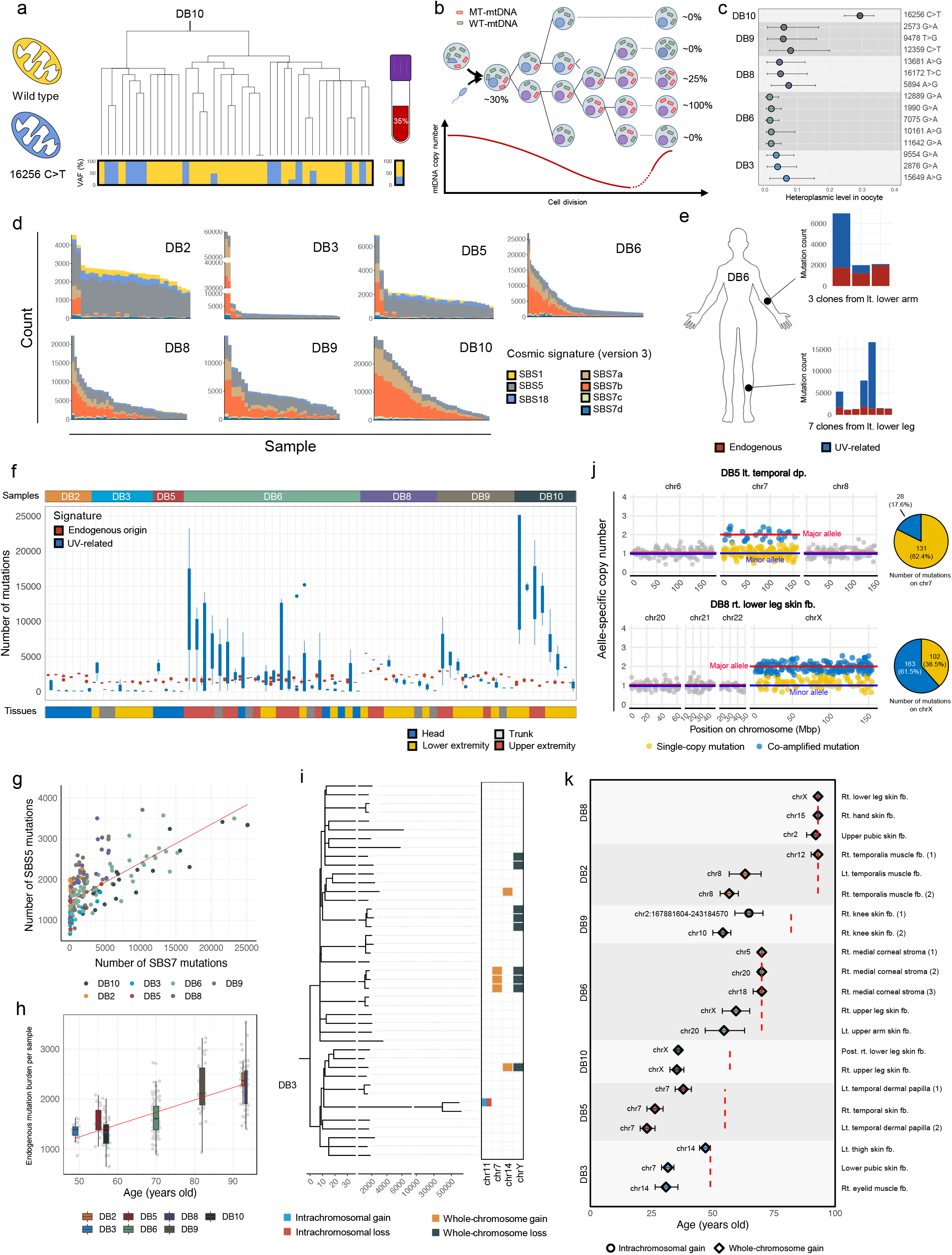
Mitochondrial DNA heteroplasmy in oocytes and late-stage mutations harbored in clones. **a,** an example of mtDNA mutation that was found in multiple clones of DB10(top) and polyclonal blood (right) in different heteroplasmic levels (bottom). **b,** a schematic representation demonstrating mitochondrial bottleneck during embryogenesis and random segregation of mtDNA during mitosis, which lead the early mtDNA variant to variant heteroplasmic levels in adult cells. **c,** mtDNA variants and their heteroplasmic levels estimated to be harbored in the first cell of the five individuals. **d,** total number and signature of somatic mutations in each of the 334 clones. **e,** examples of variable burdens of UV-related mutations among clones established from thesame anatomical locations. **f,** a massive heterogeneity of UV-mediated mutational burden among clones established in the same anatomical location (interclonal distance < ~1cm). **g,** linear correlation between the numbers of SBS7 (UV-mediated) and SBS5 (an endogenous, clock-like) mutations in skin fibroblast clones. Approximately one additional SBS5 mutation is formed per ten SBS7 mutations. **h,** the rate of purely endogenous mutations in skin fibroblasts showing a linear correlation with age (24.3 substitutions per year). **i,** large-segmental copy number changes (>10Mbps) with the developmental phylogenetic tree of DB3. **j,** examples of the gains in chromosomal copy number that occurred in the third and tenth decades of life (top and bottom, respectively). **k,** timing estimation of the copy number gains (> 50Mbps) observed in the clones. Ages at death are shown in red dashed lines. fb., fibroblast.

### Late-stage mutations and signatures

Most somatic mutations identified (overall 1,688,652 SNV and indel mutations from 334 clones established from seven donors) were not shared by multiple clones, therefore were post-embryonic, late-stage mutations accumulated over a lifetime. We first explored prevalence of cancer-associated variants, but any apparent driver mutations deposited in the catalog of cancer mutations were not identified. In addition, we did not find any evidence of positive selection from these mutations with a previously described dN/dS ratio model^37^. Mutational signature analyses suggest that the vast majority of mutations were attributable to a few known mutational processes, including UV-mediated DNA damage (SBS 7a-7d) and endogenous clock-like mutagenesis (SBS5 and 1) (**Fig. 4d**). Similar to non-malignant bronchial epithelium exposed to tobacco-smoking and hepatocytes^38,39^, there was a large variation in the number of UV-mediated mutations despite clones being established from a similar location (**Figs. 4e, 4f**). These results may be caused by the original depth of the fibroblasts in the skin tissues and/or by the heterogeneous DNA repair capabilities across the clonal lineages^40^. Two corneal keratinocyte clones, which are more directly exposed to UV, harbored an extremely high number of UV-mediated mutations (n=58,858 and 61,146 in DB3), which was comparable or higher than the mutational burden of most non-hypermutated cancers^41^. From all the clones from the seven individuals, we identified a positive correlation between the SBS 7 (UV-mediated) and SBS 5 (endogenous, clock-like) mediated mutations (adjusted R-squared = 0.75, p < 0.001; **Fig. 4g**). Given the endogenous, clock-like properties of SBS 5, it seems unlikely that the amount of SBS 5 mutations associated with SBS 7 is caused by direct damage from UV. It is more plausible that UV-mediated DNA damages activate the molecular mechanism underlying SBS5, as also suggested in tobacco smoking-mediated mutagenesis^42^. The linear correlation enabled us to determine the number of pure, endogenous mutations after adjusting for the impact of the UV-mediated damage. We estimated that approximately 24 endogenous base substitutions accumulated per year in the skin fibroblasts (**Fig. 4h**). The rate is in line with that of the human stem cells in the colon, liver, and lung^38,39,43^.

### Copy number changes and structural variations in normal cells

Finally, we investigated somatically acquired structural variations in the clones. Although the vast majority of the clones showed diploid genomes, 10% to 23% of the lineages in each individual have shown somatic copy number variations in DNA segments longer than 10Mbp. Some of these variations were shared in the close lineages (**Fig. 4i**), confirming that the event was acquired *in vivo*. Breakpoints of genomic rearrangements often exhibited clusters of base substitutions, primarily APOBEC-mediated C>T and C>G substitutions at the TpC dinucleotides (**Extended Data Fig. 7a**), suggesting single-strand DNAs are exposed in the processes of double-strand break formation and repair^44,45^.

Of the copy number variations, 70% (38 out of 54) were chromosome-level copy number changes (**Fig. 4i, Extended Data Fig. 7b**), more frequently in chromosomes X and Y (50%, 19 out of 38) (**Extended Data Fig. 7c)**. Because chromosome Y (male) and the inactive chromosome X (female) are transcriptionally, thus functionally negligible compared to autosomes, loss or gain of these chromosomes would be more tolerated. Our data support to a recent observation of frequent Y chromosome loss among blood cells in the general human population^46,47^ and the concurrent study.

Using the density of endogenous mutations in amplicons as a molecular clock, the timing of large copy number gains was estimated, as reported in cancer (**Methods**) ^48^. Briefly, amplicons occurring late in life co-amplify many substitutions accumulated in the segment (**Fig. 4j**). However, co-amplified substitutions are less frequent in amplicons occurring early in life because of the short time period for acquiring mutations before the amplifications. Therefore, the ratio of co-amplified and non-amplified mutations correlates with the timing of an amplification event. Of the 21 informative copy number gains (>50 Mb that harbor a sufficient number of mutations for statistical testing), 13 amplicons (62%) were dated to at least 10 years earlier than the time of death (**Fig. 4k**). Seven amplification events (33%) were observed to have occurred in the first four decades of life. Our data imply that large-scale copy number changes may not be a rare event in our aged somatic cells.

## Discussion

Herein, we examined whole-body mosaicism using genome sequencing of a large number of clones from the whole-body locations of individuals. For detection of somatic mutations in normal human cells, a few different approaches have been used such as whole-genome amplification^49^, laser capture microdissection^39,43,50,51^, sequencing of small biopsies that contain limited numbers of microscopic clones^52–55^, and massive clonalization of single cells^15^ as used in this study. Each approach has its pros and cons. Although the clonalization method is applicable only to dividing cells and requires a substantial experimental effort, it provides the most sensitive and precise clonal mutational calls, which is essential for reconstructing unambiguous developmental phylogenetic trees given the marginal mutation rate in each cell division. Unlike whole-genome amplification, the sensitive and specific mutation calls in the sequencing of clones also enable the investigation of the actual mutation burden in original single cells and the exploration of accurate mutational signatures that shaped the mutations in the somatic cellular lineage over a lifetime.

We established 40 - 120 clones per individual, which enabled the robust exploration of embryogenesis to the ~3rd to ~9th cell divisions depending on the number of stochastic spontaneous mutations and the contribution of an early lineage in adult tissues. Combining with WGS of blood and targeted deep sequencing of various bulk tissues, tracing of early cells into diverse human tissues was possible. Our approach showed the pervasive but stochastic unequal contribution of early cells to the entire human body. Additionally, we showed the early emergence of asymmetry to different body parts, such as the three germ layers, left-right and cranio-caudal axes, and hematopoietic tissues from the largest-scale clonal sequencing of warm-autopsies to our knowledge.

This study demonstrates that human embryogenesis can be further dissected to the later cell stages by scaling up our approach, investigating a higher number of clones, bulk tissues, and single cells from more diverse organs, tissues, and cell-types. More comprehensive developmental trees could be reconstructed by including more tissues such as extraembryonic tissues (placenta) and ectoderm-origin tissues. As the costs of WGS drop and with further technological advances in single-genome sequencing technologies, sequencing of 10,000 or more cells would be a deliverable project in the very near future that will comprehensively elucidate central questions relating to the nature of early human embryogenesis.

Although we mainly focused on common characteristics among individuals in this study, human embryogenesis is, in principle, a personal event and thus may be substantially variable across individuals. Presumably, a proportion of developmental diseases are induced by catastrophic clonal dynamics in embryogenesis. In addition, since early embryonic mutations will be harbored in a substantial fraction of somatic cells in an individual, they may contribute to a wide range of diseases depending on the functional consequences and cell types they are present in. Similar analyses of more individuals, particularly for diseases with unknown etiologies, will likely yield the clinical impact of early human embryogenesis to an unprecedented resolution in the future.

## Data availability

Whole-genome and targeted sequencing data are deposited in the European Genome-phenome Archive (EGA) with accession id: EGAS00001004824.

## Code availability

In house scripts for genomic analyses and simulation studies are available on GitHub (https://github.com/seongyeol-park/Human_Lineage_Tracing and https://github.com/chrono0707/Human_Lineage_Tracing). All other code is available from the authors on request.

## Acknowledgements

We are deeply indebted to the donors who donated cells and tissues for this study. We thank Michael R Stratton, Peter J Campbell, Tim HH Coorens, Raheleh Rahbari, Luiza Moore (Wellcome Sanger Institute) and Maksim V. Plikus (University of California, Irvine) for their fruitful comments and discussions. We also appreciate Jong-Yeon Shin (Macrogen Inc), Mee Sook Jun, Heejun Jung (Kyungpook National University), Jae Ho Lee, Hyun Su Lee (Kyemyung University), for their methodological advice and technical assistance. This work was supported by the Korea Health Technology R&D Project through the Korea Health Industry Development Institute funded by the Ministry of Health & Welfare of Korea (HI17C1836 to Y.S.J.), the Suh Kyungbae Foundation (SUHF-18010082 to Y.S.J.), a National Research Foundation (NRF) of Korea funded by the Korean Government (NRF-2020R1A3B2078973 to Y.S.J.; NRF-2020R1A5A201732 and NRF-2019R1I1A3A01060675 to J.W.O.; NRF-2020R1A6A3A01100621 to S.P.; and NRF-2019H1D3A2A02061168 to S.Y.K.).

## Author contributions

Y.S.J., J.W.O. and S.P. conceived the study; N.M.M. designed the warm autopsies and developed the entire protocol of the clonal expansion and bulk tissue preparation with help from J.W.O.; J.W.O., N.M.M., J-W.C, J.M.P., D.K., J.H.C., S.G.K., J.H.H., M.K., D.S.K., J.Y.P., T.K., J.Y., and Y.A. conducted autopsies, tissue sampling and clonal expansions. S.P. and R.K. conducted most of the genome and statistical analyses with a contribution from S.Y.K. and Y.S.J.; Ju.L. and J.W.P. contributed to large-scale genome data management. Jo.L. conducted mutational signature analysis. Y.S.J., S.P., R.K., N.M.M and J.W.O. wrote the manuscript with contributions from all the others. Y.S.J. and J.W.O. supervised the study.

## Extended Data Figure legends

**Extended Data Fig 1.**
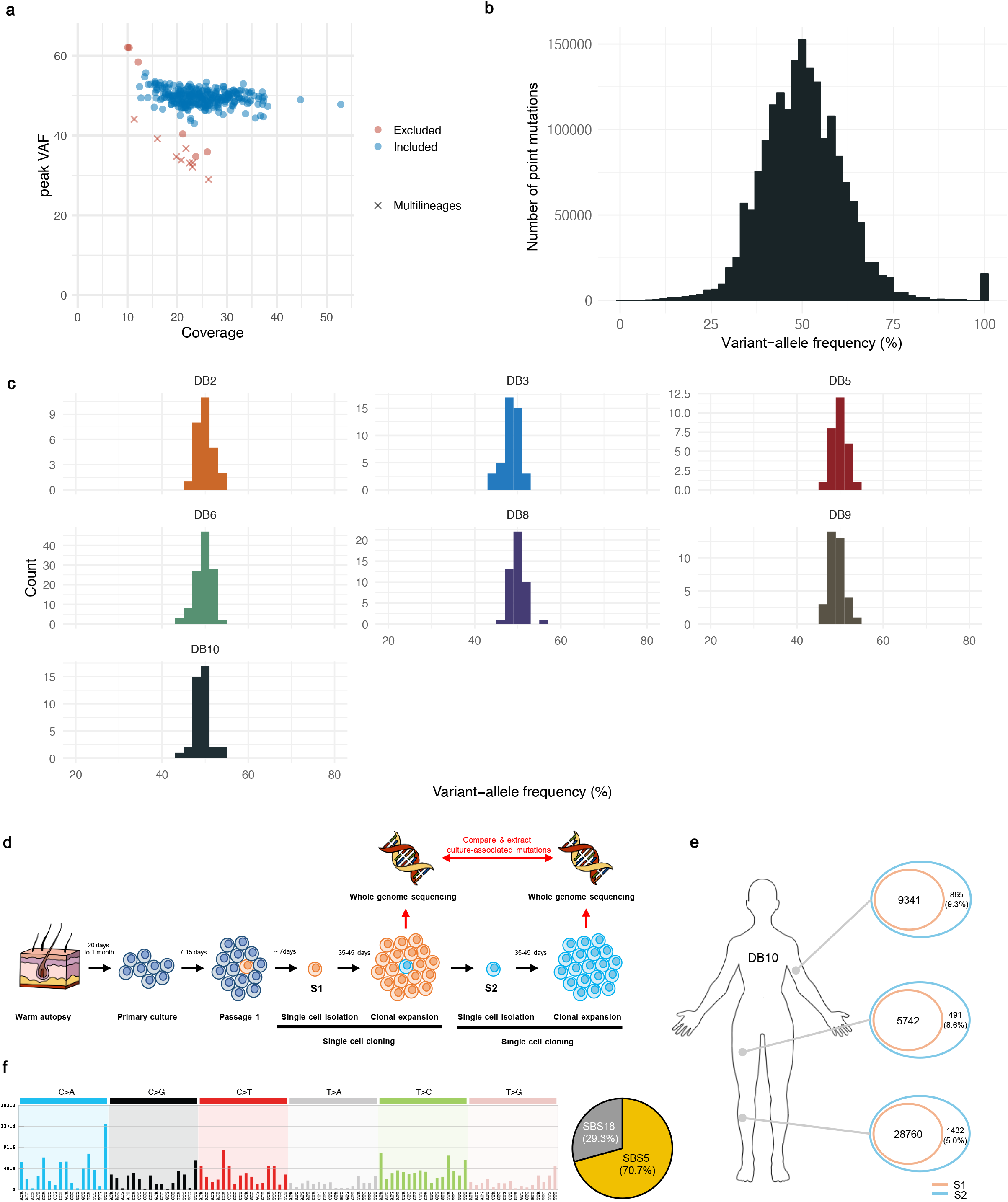
Quality assessment of single-cell-derived clones. **a,** a scatter plot showing peak VAF and mean coverage of 374 clones. X-shaped dots represent colonies established from two or more single-cells. Single-cell driven colonies (clones) with peak VAF near 50% were included in this study. **b,** a histogram demonstrating the distribution of VAFs of all somatic mutations detected in the study (*n* = 1,688,652). **c,** histograms demonstrating the distribution of peak VAFs of clones per individual. **d,** the experimental design for detecting culture-associated mutations. We isolated a single cell (S1) from the primary culture of a warm autopsy sample and another single cell (S2) from the S1-derived clone. Following clonal expansion for 35-45 days, we sequenced S1- and S2-derived clones and extracted culture-associated mutations by comparing them. **e,** Number of mutations in S1 and S2 from three warm autopsy samples of DB10. The clonal mutations in S2 which are not clonal in S1 were considered to be culture-associated. **f,** the signatures of the culture-associated mutations which are delineated by COSMIC SBS5 (70.7%) and SBS18 (29.3%).

**Extended Data Fig 2.**
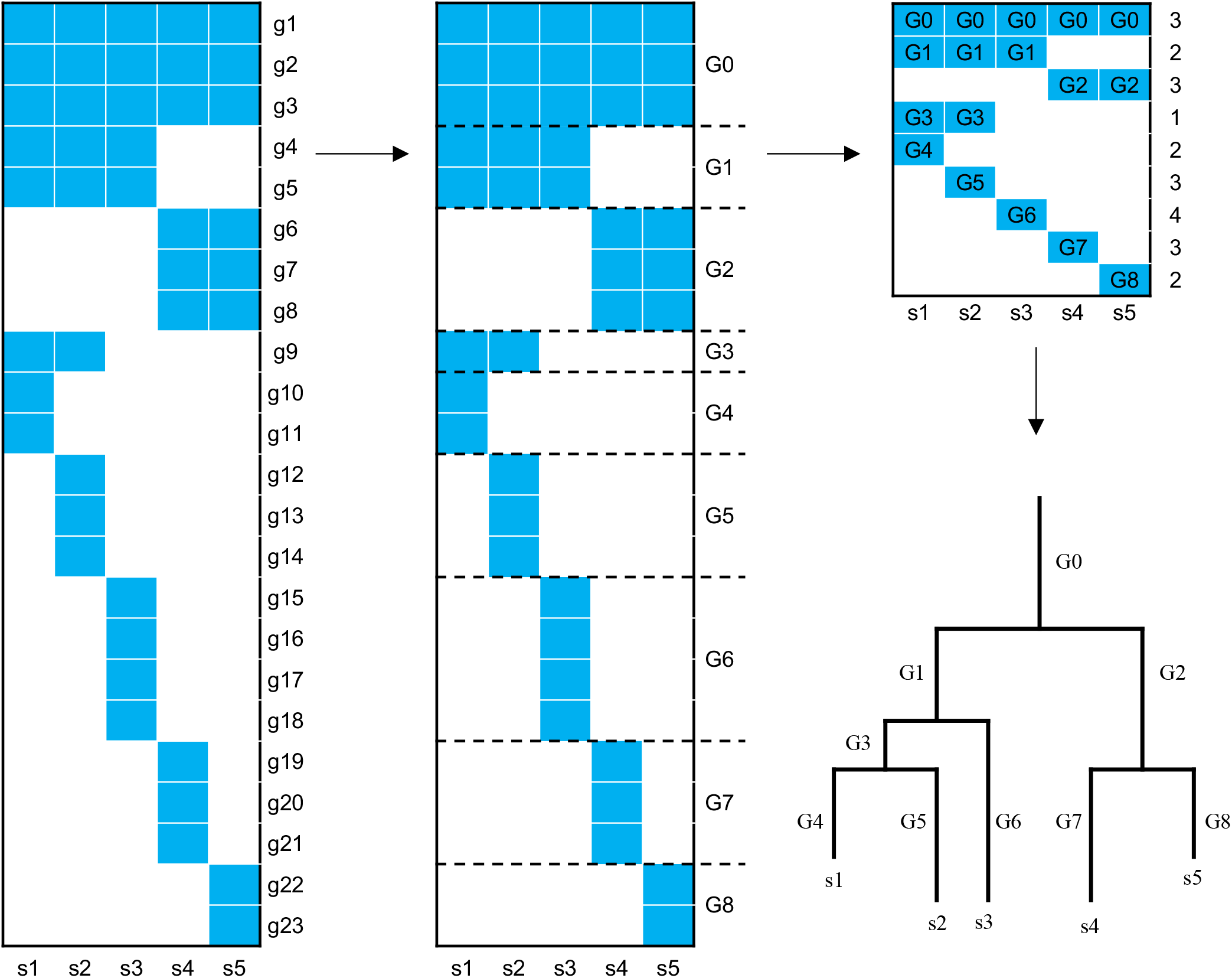
Graphical method summary. A schematic illustration demonstrating our approach reconstructing a developmental phylogenetic tree. Let S= [s_1_, s_2_, …, s_5_] be the set of 5 clones (samples), and G= [g_1_, g_2_, …, g_23_] be the union set of mutations detected in one or more samples from the same individual. We then build a matrix M with rows labeled g_1_, g_2_, …, g_23_, and columns labeled s_1_, s_2_, …, s_5_. If the VAF of somatic mutation g_i_ in sample s_j_ is greater or equal to a given threshold, M_ij_ was assigned to 1 (blue-colored tile), while others to 0 (white-colored tile). After removing germline variants, we grouped all mutations with the same profile into a mutation group according to the sharing pattern between samples. Over 8 distinct mutation groups in this example, mutation matrix M_8×5_ is defined such that each column represents a sample and each row represents a mutation group. From the mutation matrix M_8×5_, we reconstructed a phylogenetic tree.

**Extended Data Fig 3.**
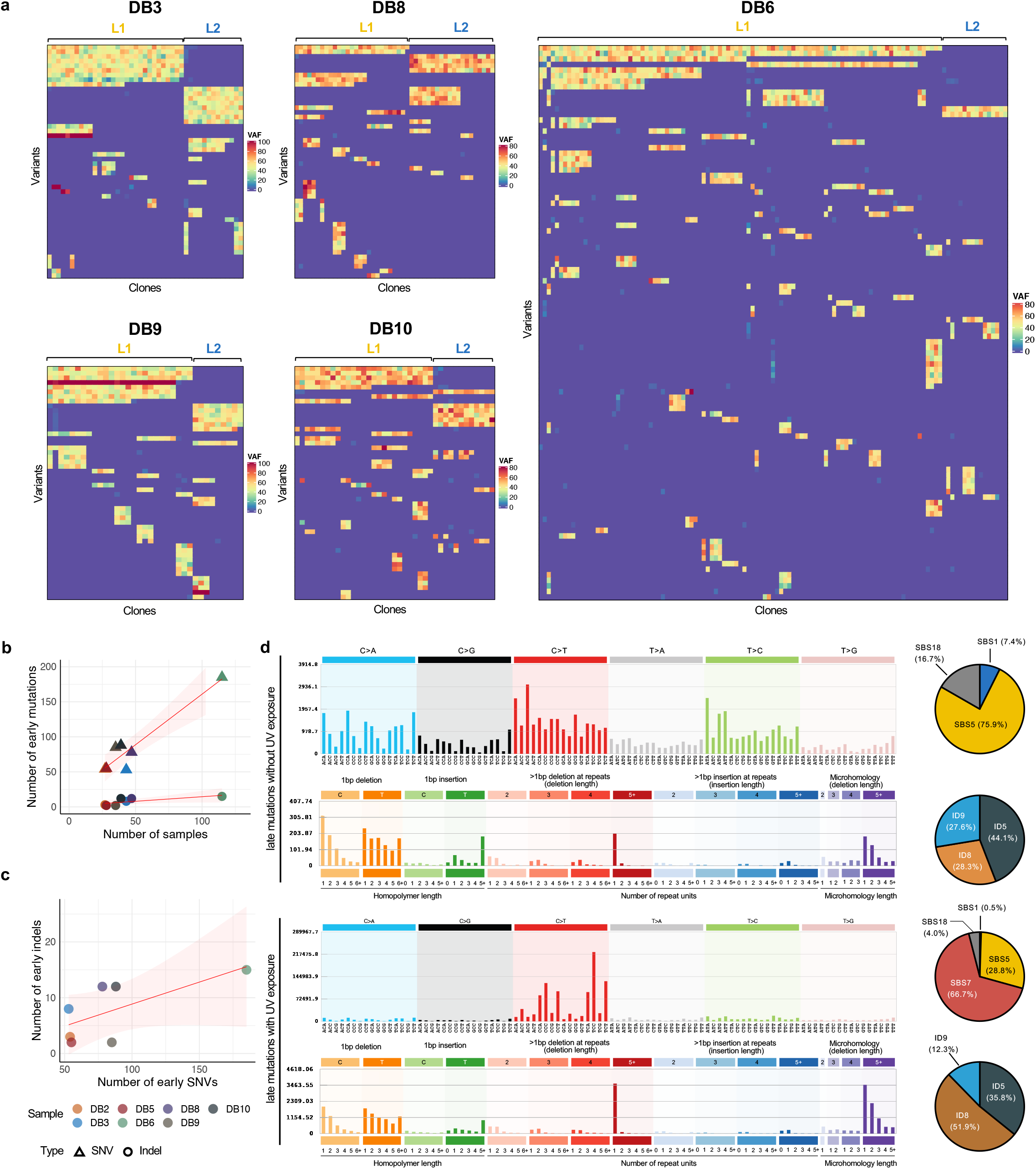
Somatic mutations identified from the single-cell derived clones. **a,** heatmaps illustrating the VAFs of mutations in single-cell-derived clones. The earliest 100 mutations were illustrated in DB6. From the other donors, the earliest 50 mutations were shown. ‘L1’ denotes the major lineage in the initial branching of the phylogenetic tree, while ‘L2’ represents the minor lineage. **b,** the correlation between the number of early mutations and clones, separately calculated for single nucleotide variants (SNVs; triangle) and short indels (circle). **c,** the correlation between the number of early SNVs and indels. **d,** the mutational spectrums of late mutations. We categorized lineages into two groups according to the proportion of COSMIC SBS7 in late mutations with a cutoff of 5%. The upper panel displays the mutational spectrums of substitutions (n=74,824) and indels (n=3,404) in clones with SBS7 >5% (with UV exposure), while the lower panel displays the spectrums of substitutions (n=1,457,489) and indels (n=31,805) in clones with SBS7 ≤5% (without UV exposure).

**Extended Data Fig 4.**
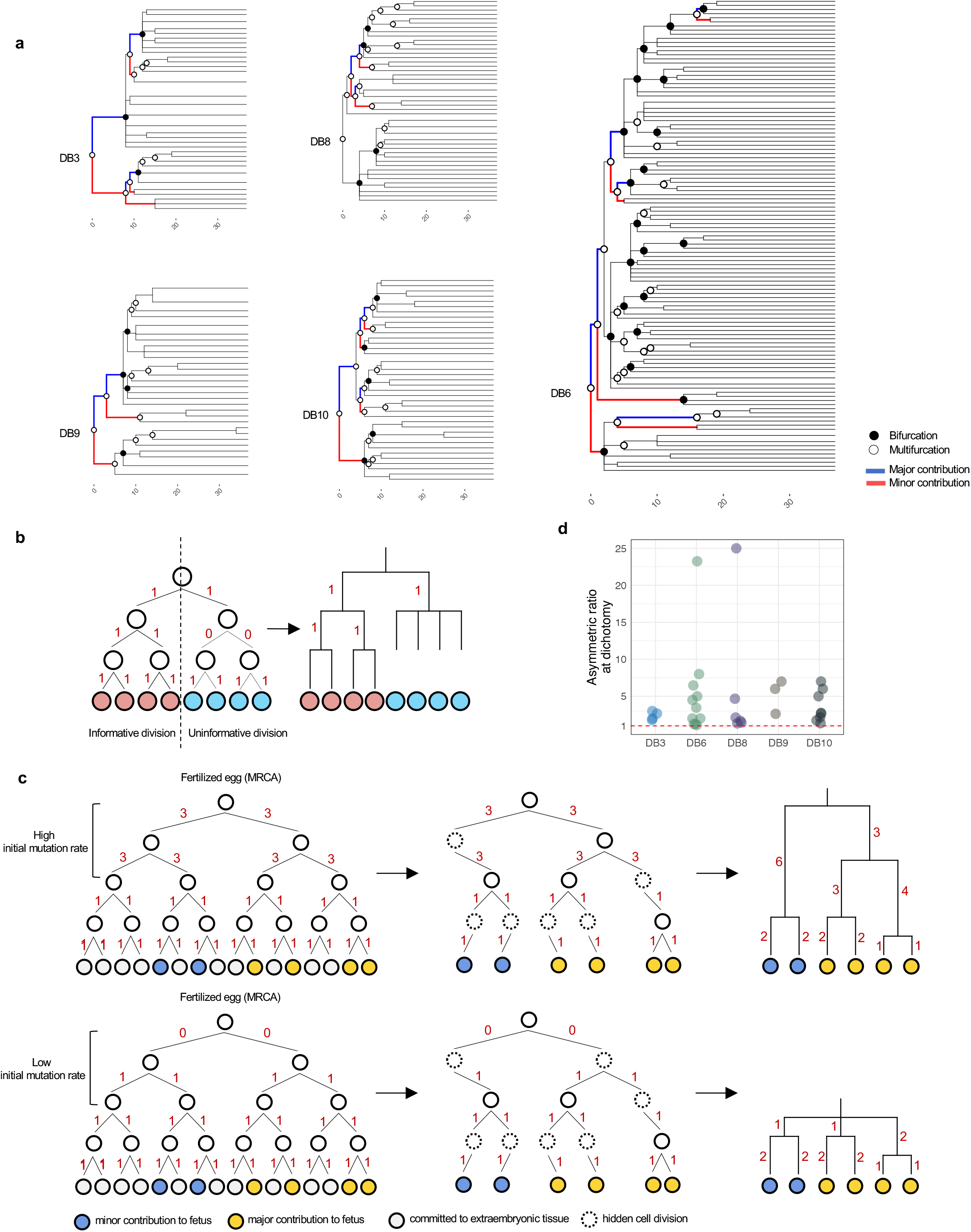
The structure of phylogenetic trees. **a,** annotated phylogenetic trees of DB3, DB6, DB8, DB9, and DB10. Dichotomy (bifurcation) and polychotomy (multifurcation) nodes are indicated by black-filled circles and hollow circles, respectively. At each bifurcation node, two lineages were colored depending on the degree to which they contribute to subsequent descendant lineages: the major contribution lineage with red, and the minor contribution lineage with blue. **b,** a schematic illustration demonstrating informative and uninformative cell divisions. In contrast to a cell division accompanying spontaneous mutations (informative division, left), a cell division without intrinsic mutation cannot be reflected in a phylogenetic tree (uninformative division, right). **c,** a schematic illustration showing the effect of initial mutation rate on the number of first lineages. **d,** the asymmetric ratio at bifurcating nodes, which is defined as the number of descendants of the major contribution lineage divided by that of minor contribution lineage.

**Extended Data Fig 5.**
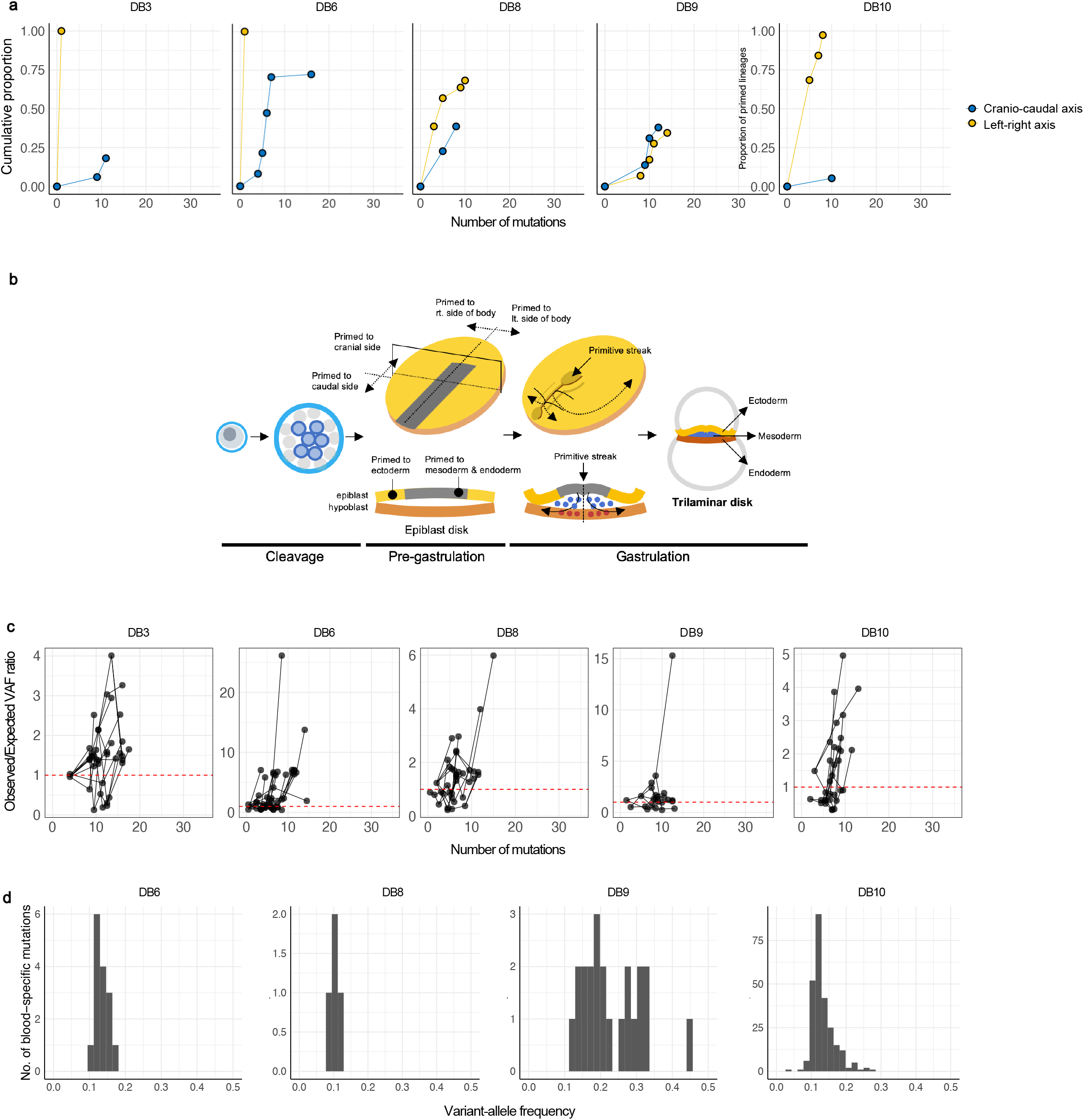
Early cell dynamics constituting various asymmetries. **a,** cumulative proportion of lineages with asymmetric distribution for the axis of interest (left-right or cranio-caudal) over the course of molecular time of mutations. The number of mutations of each dot is the mean value from a subgroup of mutations in each branch (**Methods**). **b,** a schematic illustration showing the emergence of primitive streak and formation of axes in early embryogenesis. **c,** ratios of VAF observed in bulk blood to expected VAF from phylogenetic trees. The number of mutations of each dot is the mean value from a subgroup of mutations in each branch (**Methods**). Dots in the same phylogenetic lineage were linked by lines. **d,** Histograms of VAFs for blood-specific mutations.

**Extended Data Fig 6.**
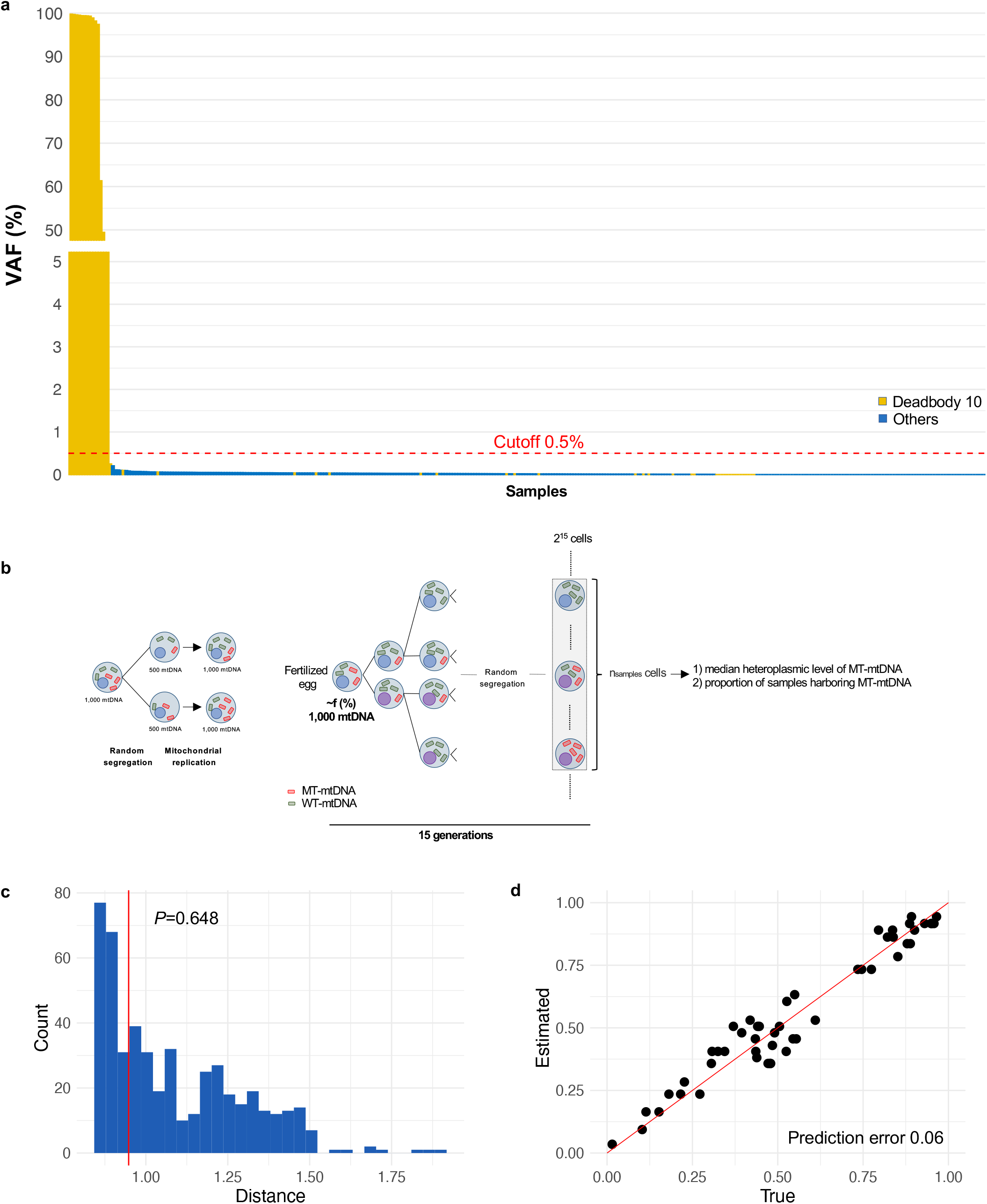
Heteroplasmic mitochondrial variants in a fertilized egg. **a,** a variant allele frequency (VAF) of MT:16,256 C>T substitution in samples from DB10 and others. Applying a VAF cutoff of 0.5%, the variant was found only in the 14 clones establishedfrom DB10 but not in other cases. **b,** a developmental model for inferring the heteroplasmic level of a mitochondrial variant in a fertilized egg. We assumed that *f*% of mtDNA in a fertilized egg has a functionally neutral mtDNA variant (MT-mtDNA), which randomly segrate to daughter cells in successive cell divisions. Two summary statistics were drawn from this model: 1) the proportion of samples harboring MT-mtDNA (*p*), the median heteroplasmic level of MT-mtDNA (*h*). We compared the summary statistics (*p, h*) of each simulation to the observed summary statistics, and constructed the posterior distribution of *f* using the neural network regression algorithm of an approximate Bayesian computation. For detail, see **Methods** section. **c,** a histogram of the null distribution of the statistic for goodness of fit test assuming our model. **d,** Cross-validation to access the accuracy of parameter inference.

**Extended Data Fig 7.**
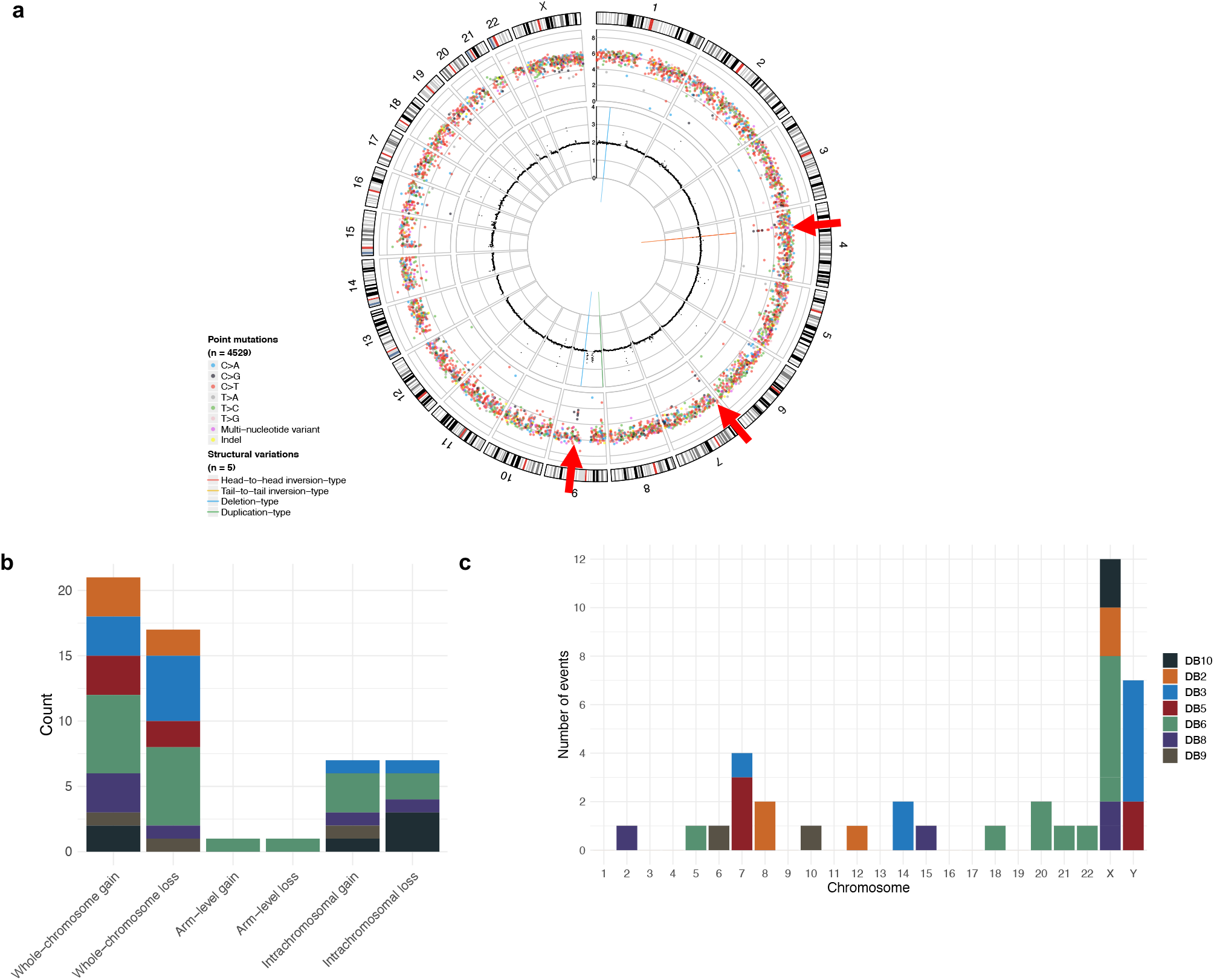
Structural variations and copy number variations in normal cells. **a,** A circos plot showing an example of kataegis accompanied by structural variation in a normal cell. From outer to inner space, log10 intermutation distance (bps), absolute copy number, and structural variations are described. Kataegis are marked by red arrows. **b,** Bar plot showing the number of copy number variations (> 50Mbps) detected in normal cells. **c,** Bar plot showing the number of copy number variations per chromosome.

## Supplementary Tables

**Supplementary Table 1. Demographic features and sample information of seven individuals.**

**Supplementary Table 2. Cell types, tissue locations, and mutational features of 334 single-cell clones.**

**Supplementary Table 3. Tissue types and locations of 374 bulk tissues recapturing early mutations via targeted deep sequencing.**

**Supplementary Table 4. Phylogenetic structure and copy number variations of all lineages in seven individuals.**

## Methods

### Warm-autopsy and tissue sampling

A total of seven donors transitioning to imminent death were included in post-mortem tissue donation for this study (**Supplementary Table 1**). The objectives and procedure of this study were explained to the patients or nearest kin, and permission for autopsy was obtained before their death. From June 2016 to January 2019, ten immediate autopsies had been performed at the Kyungpook National University Hospital and Department of Anatomy at Kyungpook National University, School of Medicine. In seven of them primary culture of tissues was successful. When a participant died, we collected 20-30 ml of blood in a purple-topped, heparinized tube. The whole body was sterilized by 70% ethanol before tissue sample collection. The skins, subcutaneous tissues, muscles, and hair follicles from several randomly chosen anatomical sites and corneas were taken and immersed in transportation media as soon as possible (**Fig. 1b**). The internal organs were exposed through an anterior Y-shaped incision extending from the shoulders to the pubis and dissected from their posterior attachments to the body. Tissues were collected as fresh tissue, snap-frozen tissue using liquid nitrogen (stored at −70°C), and formalin-fixed tissue. Fresh tissues were transported to the laboratory for tissue culture experiments. All tissues were collected within 36 hours of death and processed on the day of the collection according to the protocol. All the procedures in this study were approved by Kyungpook National University Institutional Review Board and Human Research Ethics Committee (approval number: KNU-2019-0088 and KNU-2019-0151). This study was conducted in accordance with the Declaration of Helsinki provisions. No statistical methods were used to predetermine the sample size.

### Primary culture of tissues

We obtained a total of 165 tissues, approximately 1 × 1 × 1 cm^3^ in size, for a single-cell expansion experiment from the seven autopsy cases (**Fig. 1b, Supplementary Table 2**). All tissues were processed as previously described with minor modifications. Growth medium contained Dulbecco’s modified Eagle’s medium (DMEM) low glucose, 20% fetal bovine serum, 100 IU/ml penicillin, 100 μg/ml streptomycin, 2mM L-glutamine and 1 μg/ml fungizone. All components were obtained from Gibco (Thermo Fisher Scientific, Waltham, MA, USA). The detailed protocols for each tissue type are described below.

### Skin culture

The dermal skin fibroblasts were cultured by the explant method described previously with minor modification^57–59^. Briefly, the skin samples were washed 2-3 times with buffer saline (1X PBS; Gibco, Thermo Fisher Scientific). The adipose tissue and associated blood vessels were removed. Subsequently, the tissues were cut into 1-2 mm^2^ pieces and put in 1mg/ml collagenase and dispase (C/D) solution (Roche Diagnostics GmbH Roche Applied Science, Mannheim, Germany) at 37°C for 1 hour. After the treatment, the epidermis layer was separated from the dermis layer. The dermis tissue was washed by 20% FBS (Gibco, Thermo Fisher Scientific) containing media to inhibit C/D activity. Then, the dermis tissue was minced into sample pieces and cultured in collagen I coated 24 well plates (Corning BioCoat™, NY, USA) with 200 μl media in a humidified incubator at 37°C and 5% CO_2_ in the air.

### Muscle culture

The skeletal muscle and other smooth muscle were used for muscle culture with a previously described protocol with minor modification^60^. The muscle tissue was washed by 1X PBS (Gibco, Thermo Fisher Scientific) and then FBS containing medium. Next, the connective tissue and adipose tissues were removed. The tissue was chopped into small pieces of 1-2 mm^2^ and digested for 45-60 minutes at 37°C with 1mg/ml C/D solution with frequent agitation. The tissue slurry was washed with FBS containing growth medium and centrifuged at 1400 rpm for 5 minutes at room temperature. The pellet was resuspended in a fresh 20% FBS containing growth medium. A small piece of tissue was taken out from a slurry and minced with sharp blades and cultured in collagen I-coated 24 well plates with 200 μl media in a humidified incubator at 37°C and 5% CO_2_ in the air.

### Corneal tissue culture

The corneal tissue was washed by 1X PBS (Gibco, Thermo Fisher Scientific). Then, the cornea was cut diagonally into 16 pieces, and each piece was transferred into each well of collagen I coated 24 well plates. The fragments of cornea tissue were chopped in a culture plate with 20% FBS containing DMEM medium. For cornea keratinocyte, the medium was replaced by keratinocyte specific CNT-PR medium (CELLnTEC Advanced cell system, Switzerland) after 24 hours of tissue culture.

### Hair follicle culture

Hair follicle samples were collected from the scalp and other anatomical sites of individuals. Excess hair shafts were trimmed with scissors and washed by 1X PBS. The single hair follicles were isolated and cultured using the previously described methods after the determination of hair follicle stages^61^. The anagen hair follicles were used for dermal papilla (DP) and dermal sheath (DS) culture. DP and DS were isolated from the bulbs of micro-dissected hair follicles. Each DP and DS were separately transferred to collagen I coated culture dishes and culture with 20% FBS containing growth medium.

### Cell outgrowth and passage

The medium was changed every four days after culture to remove cellular debris in suspension. Usually, the cells were outgrowth within two weeks, depending on tissue and cell types. After sub-confluent, the cells were maintained in DMEM supplement with 10% FBS at 60 mm or 100 mm dishes to adjust the microenvironment of cell growth.

### Single-cell cloning and clonal expansion

The sub-confluent cells were washed by ice-cold 1X PBS and harvested by using 1X trypsin (TrypLE™, Gibco, Thermo Fisher Scientific) in 15ml conical tube (BD Falcon, USA) with sufficient amount of fresh medium to neutralize the trypsin activity. The cell pellet was resuspended by 1ml growth medium, and the total cell number was counted by trypan blue using a disposable hematocytometer slide in an automatic cell counter (EVE™ automatic cell counter, NanoEnTek, Inc., USA). After the calculation of the total cell number, a serial dilution method was applied to dilute the cells up to 10 cells/ml. Then, 100μl cell suspension medium was seeded in each well of 96 well plates (BD Biosciences, Franklin Lakes, NJ, USA) and incubated at 37°C with 5% CO_2_. Each well contains a single cell. Within 12-15 hours, the cell was manually checked under a light microscope (Olympus, Tokyo, Japan) at 10 × magnification, and the wells containing a single cell were selected. Single cells were proliferated up to 80-90% confluency; afterward, the individual cell clones were expanded into 24 well plates. The confluent clones were collected for DNA extraction and whole-genome sequencing.

### Library preparation and whole-genome sequencing

We extracted genomic DNA materials from clonally expanded cells and matched peripheral blood using DNeasy Blood and Tissue kits (Qiagen, Venlo, Netherlands) according to the protocol. DNA libraries for WGS were generated by an Accel-NGS 2S Plus DNA Library Kit (Swift Biosciences, Ann Arbor, MI, USA) from 1 μg of genomic DNA materials. WGS was performed on either the Illumina HiSeq X platform or the NovaSeq 6000 platform to generate mean coverage of 25.2X for 374 clonally expanded cells and 94.8X for 7 matched blood tissues.

### Variant calling and filtering of whole-genome sequencing data

Sequenced reads were mapped to the human reference genome (GRCh37) using the BWA-MEM algorithm^62^. The duplicated reads were removed by Picard (available at http://broadinstitute.github.io/picard), and indel realignment and base quality score recalibration were performed by GATK^63^. Initially, we identified base substitution and short indels by using HaplotypeCaller^64^ and VarScan2^65^. Variants called by both of the tools were included for future analysis. For every variant in any of the single-cell derived clones, we evaluated the number of mutant and wild type reads. To establish the high-confident somatic variant sets, we applied additional filtering processes for these variants as follows:

1. The sequencing and mapping artifacts of the initially identified variants were filtered out by using a locus-specific background error matrix, which was generated by in-house normal tissue WGS datasets (panel of normal).
2. If a variant had supporting reads in all samples from an individual, we considered it as germline variants and filtered it out. The variants, which are absent only in samples with loss of heterozygosity, were also considered as germline variants.
3. Variants with the following condition were considered to be sequencing or mapping artifacts and removed.

⍰ Variants with less than three supporting reads.
⍰ The difference in median mapping quality between reference and variant reads is greater than 10.
⍰ The median mapping quality of reference reads is less than 40.
⍰ The median mapping quality of mutant reads is less than 40.
⍰ The median number of mismatched bases in mutant reads is greater than five.
⍰ If more than 90 percent of supporting reads were soft-clipped or had additional insertion/deletion.
⍰ If five or more samples have only one or two variant reads.

We visually inspected the variants passed the filtering process using the Integrative Genomics Viewer (IGV) ^66^.

### Quality control of samples

Among a total of 374 clonally expanded cells, 18 with low depth of coverage (mean depth < 10) were excluded. Based on the VAFs of somatic mutations and established phylogenetic trees, we removed additional 19 samples thought to be multiclonal: they have variants in mutually exclusive lineages simultaneously, and/or low peak VAFs (**Extended Data Fig. 1a**). In addition, three samples which show unexplainable high peak VAFs were excluded for data integrity. Finally, we analyzed the remaining 334 whole-genome sequencing data from single-cell derived clones to reconstruct phylogenetic trees.

### Detection of culture-associated mutations

Since we performed two times of *in vitro* culture, culture-associated mutations could be integrated in our mutation list. First, the mutations which were generated in short-term culture before single cell isolation would exist in all cells in clones (clonal). Second, the mutations which occurred after single cell isolation would exist in part of cells in clones (subclonal). The latter mutations seem to be rare, because additional VAF peaks < 50% were not detected in most 334 single-cell derived clones and in the merged VAF distribution of all somatic mutations (n = 1,688,652; **Extended Data Fig. 1b, c**). To approximate the amount of the culture-associated mutations before single cell isolation, we conducted serial single-cell clonal expansions followed by WGS of both clones in three samples (**Extended Data Fig. 1d**). In these experiments, mutations, which are clonal in second clones but subclonal or not detected in first clones, are culture-associated mutations that occurred during first single cell cloning. To discriminate clonal mutations and subclonal mutations, we calculated mutant-cell fraction (MCF) by the same method estimating cancer cell fraction in the previous studies^48,67^. Mutations, which showed MCF < 0.25 in first clone and MCF > 0.25 in second clone simultaneously, were defined as culture-associated mutations.

### Reconstruction of the phylogenetic trees and defining early mutations

The developmental phylogenetic tree of an individual was inferred from multi-sample genotype information. We assumed that the chance of recurrent somatic mutations in multiple lineages are negligible. Let S=[*s*_1_, *s*_2_, …, *s*_*n*_] be the set of n samples, and G=[*g*_1_, *g*_2_, …, *g*_*m*_] be the union set of mutations detected in one or more samples from the same individual. We then build a matrix M with rows labeled *g*_1_, *g*_2_, …, *g*_*m*_, and columns labeled *s*_1_, *s*_2_, …, *s*_*n*_, If the VAF of somatic mutation *g*_*i*_ in sample *s*_*j*_ is greater or equal to a given threshold *M*_*ij*_ was assigned to 1, while others to 0. Mutations shared in all samples were considered to be germline variants. After removing these germline variants, we grouped all mutations with the same profile into a mutation group according to the sharing pattern between samples. (**Extended Data Fig. 2**) Over *m′* distinct mutation groups, mutation matrix *M*_*m′×n*_ is defined such that each column represents a sample and each row represents a mutation group. From the mutation matrix *M*_*m′n*_, we reconstructed a phylogenetic tree by using graphic theoretic methods^68^. Mutations shared by two or more samples, which cannot be rooted to a most recent common ancestor other than fertilized egg, were considered to occur independently in those samples, and therefore were not employed to reconstruct a phylogenetic tree. A phylogenetic tree for the mutation matrix *M*_*m′n*_ is a rooted tree where each sample is attached to exactly one terminal node of the tree, and each of the *m′* mutation group is associated with exactly one branch of the tree, with the number of mutations in the corresponding mutation group being the length of the branch.

Reconstructed phylogenetic trees were manually converted into Newick format. The trees were drawn in R using ‘ggtree’ package^69^. Most samples shared the small number of mutations (< 35) except for some samples which shared more than hundreds of mutations. Only the mutations shared in small amount (<35) among samples were defined as early mutations. To validate the early mutations, five biological replicates (culturing the sample twice from three samples, and separating cultured cell pool into two from two samples) and two technical replicates (sequencing the DNA pool twice) were made. All the detected early mutations were validated in the replicates.

### Estimating the number of inner cell mass precursor cells and mutation rate in early embryogenesis

Several studies implicate the presence of a developmental bias among blastomeres at an early stage of embryogenesis^70–74^. However, little is known regarding to the exact details of this critical event in human embryogenesis, and both the precise timing and number of cells contributing to epiblasts have yet to be clarified. We have attempted to answer these parameters, using the framework of an approximate Bayesian computation (ABC) ^75^. Our simplified model of early human embryogenesis has four parameters: the number of blastomeres contributing to epiblasts (*n*), the stage at which the precursors of epiblasts are chosen (*s*), the mutation rate before ZGA (*R*_≤2_), and the mutation rate after ZGA (*R*_>2_) (**Fig. 2c**). ZGA was assumed to occur at the 4 cell stage of early embryogenesis^23,25^. From a fertilized egg, cells are doubled at each stage, and somatic mutations accumulate in the nuclear genome with mutation rate *R*_≤2_ before ZGA and *R*_>2_ after ZGA. At a stage *s*, *n*_*preEPI*_ cells are committed towards precursor of epiblasts, and the cells divide 15 more times forming *n*_*preEPI*_ × 2^15^ cells. Of these simulated cells, *n*_*samples*_ cells (the number of clones established in each individual) are randomly selected for reconstructing a phylogenetic tree by using the somatic mutations accumulated in each cell. The major assumptions of this model are as follows:

1. All the cells in an embryo are simultaneously doubled at a stage. (that is, all the cells have the same division rate)
2. All the cells at a stage are equally likely to be epiblast precursors.
3. All the cells randomly distribute over embryo before precursor selection.
4. ZGA begins after the four-cell stage.
5. Both *R*_≤2_ and *R*_>2_ are constant during the simulation and follow the Poisson distribution.
6. Cell losses do not occur other than selection of *n*_*preEPI*_ and *n*_*samples*_ during the simulation.

This model was simulated 500,000 times for each cadaver, varying the parameters drawn from a uniform distribution with the following ranges: [2, 7] for *n*_*preEPI*_ ; 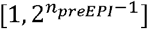 for *s*; [2.0, 10.0] for *R*_≤2_; 0.5, 2.0 for *R*_>2_. For each simulation and observed data from the five phylogenetic trees (DB3, DB6, DB8, DB9, DB10), the following set of summary statistics was extracted.

1. The number of shared mutations between more than two samples. (∑_*braches*_ *m*, where *m* represents the number of mutations assigned to each branch)
2. The number of sample groups split right after fertilization. (*n*_1*st lineage*_)
3. The number of mutations assigned to each first branch.
4. Multifurcation score defined as 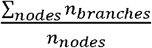, where *n*_*branches*_ and *n*_*nodes*_ denote the number of branches for each node and the total number of nodes, respectively.

The summary statistics of each simulation are compared to the observed summary statistics. The parameters of simulations that produce summary statistics that are highly similar to the observed statistics are selected. To estimate the posterior distribution of the *R*_≤2_ and *R*_>2_., we used the neural network regression algorithm implemented in the ‘abc’ package in R^76^.

### Separation of skin tissues into epidermis and dermis

We removed adipose tissue and blood vessels from the skin tissue before treatment. After washing with 1x PBS 2-3 times, we cut the tissue into small pieces. The section was placed into a C/D solution (1mg/ml) such that the epidermis layer is at the bottom. The tissue should be well covered with a C/D solution and incubated at 4°C overnight. Then, we took out the plate and peeled off the epidermis from the dermis using a pair of fine forceps. Epidermis and dermis tissues were transferred into the fresh medium and washed by 1X PBS.

### Targeted deep sequencing for bulk tissues

To estimate the contribution of early embryogenic cell lineages in adult tissues, we performed deep targeted sequencing on bulk tissues from various organs of the five individuals. Whenever possible, skin tissues were separated into the epidermis and dermis. Custom DNA bait sets were designed to include early embryonic substitutions, which were confirmed before designing the baits, and several randomly-chosen germline mutations according to the manufacturer’s guidelines (SureSelectXT Custom DNA Target Enrichment Probes, Agilent, Santa Clara, CA, USA). Of the 441 early embryonic mutations targeted, 409 mutations could have high-quality baits designed for them. DNA libraries were prepared by SureSelect^XT^ Library Prep Kit (Agilent), hybridized to the appropriate capture panel, multiplexed on flow cells, and subjected to paired-end sequencing (150-bp reads) on the NovaSeq 6000 platform (Illumina, San Diego, CA, USA) with a mean 911x depth of coverage for the early mutations. Sequence reads were trimmed and mapped to the human reference genome (GRCh37) using the BWA-MEM algorithm^62^. Reads with mapping quality ≥ 20 and base quality ≥ 20 were included for analysis. Read counts of all mutated positions in the bait-set were counted with a custom script. We only used the mutations with the sufficient read depth (> 100) in the downstream analyses.

### Subgrouping of early mutations in each branch using data from bulk tissues

Selection processes such as epiblast selection and random sampling of single cells can integrate multiple divisions into a branch in phylogenetic trees (**Fig 2d**). If a missing branch contributed to the embryo, the mutations that occurred before the hidden branching would have higher VAFs in bulk tissues than the other mutations in the same branch that occurred later. We performed the paired t-test on all possible pairs of early embryonic mutations in each branch. Mutations without significant differences in the test were clustered into a subgroup.

### Analysis of asymmetry on germ layers and anatomical axes

To investigate the distribution of early embryonic cells in each germ layer, we classified bulk tissues into several germ layer groups: ectoderm, mesoderm, endoderm, and mixed (**Supplementary Table 3**). Fresh frozen skin tissues were physically separated into the epidermis and the dermis, which were considered as ectodermal and mesodermal origins. VAFs of early mutations were compared among these germ layer groups using the Wilcox test.

We also classified bulk tissues according to the anatomical axes: left vs. right (left-right axis) or upper vs. lower (cranio-caudal axis). Tissues with the specified anatomical locations such as skin, muscle, and fat tissues were utilized in the analysis. A mid-sagittal plane and a horizontal plane crossing the xiphoid process were adopted for distinguishing left-right and cranio-caudal axes, respectively.

To compare the timing of forming asymmetry between the left-right and cranio-caudal axes, we used two approaches. First, we divided early mutations of DB6 into three groups using two thresholds of median VAF of 5% and 0.5% computed from all bulk tissues. Since VAFs of early mutations decrease rapidly as cells undergo divisions, early mutations generally show higher VAFs than late mutations. Thus, early mutations with VAF > 5% indicate the intrinsic barcodes of the earliest cells. We also inferred the cellular stages of each group using the mean mutation rate of DB6 (R≤2 = 1.37 and R>2 = 1.11) acquired from the simulation study. Using the VAF by sample matrix of each group, we performed tSNE analysis using the ‘Rtsne’ package in R^77^. In the analysis, we could identify the presence of clusters of anatomical regions and their relative timing of emergence **(Fig. 3d)**.

Second, we plotted the cumulative proportion of early lineages which had undergone significant asymmetry in the Wilcox test on the molecular time of mutations. In DB3, for example, asymmetry on the left-right axis was observed in first-occurred mutations in initial branches (**Fig. 3c**). It means that 100% of early cells underwent asymmetry at the early molecular time. By overlapping plots of the left-right and cranio-caudal axes, we compared the relative timing of asymmetry on the axes (**Extended Data Fig. 5a**).

### Tracing the dominant lineage of blood precursor cells

By calculating VAFs of early mutations in high-depth WGS of blood, we estimated the contribution of early cells to the blood cell pool. Since we observed the correlation between lineage proportion in phylogenetic trees and cellular proportion in bulk tissues (**Fig. 2b**), we compared blood VAFs (observed VAF) to the expected VAFs from phylogenetic trees. The observed/expected VAF ratio was plotted by molecular time of mutations in subgroups of mutations in each branch (**Extended Data Fig. 5c**). The molecular time of the peak ratio was defined as the emergence time of dominant lineage contributing hematopoietic tissues.

We also extracted blood-specific mutations to identify the single-cell-derived subclones and their timing of emergence. Variants were called by HaplotypeCaller^64^ and included if satisfied all of the criteria: median mapping quality of variant reads > 25, the distance of median mapping quality between reference reads and variant reads < 10, minimum mapping quality of reference reads > 0, mean base quality of variant reads > 20, the number of variant reads > 2, the proportion of clipped reads < 0.95, the median number of mismatched bases in variant reads <5, the number of single-cell clones having the same variant read < 3. All the mutations constituting the phylogenetic tree of each case were also removed. Thus, the number of final blood-specific mutations reflects the molecular time after diverging from the phylogenetic tree.

### Discovery of variants in mitochondrial genome

Because the reads sequenced from an inserted mitochondrial sequence in nuclear DNA (the nuclear mtDNA transfer, NUMT) can be misaligned to mitochondrial reference genome (chrM), we included reads mapped to chrM only if both paired reads (i) mapped to chrM, (ii) mapped properly in pair, (iii) had read length ≥ 100 bp, and (iv) had no chimeric alignment.

We filtered mitochondrial variants according to the following criteria:

⍰ We assigned a VAF cutoff to each mitochondrial variant by considering the VAF distribution locus-by-locus over all the samples. A VAF cutoff of 0.5% appeared to be optimal for the majority of variants (**Extended Data Fig 6a**). For each sample, variants with VAF lower than the cutoff were discarded.
⍰ The average variant position in supporting reads relative to read length should be between 10% to 90%. Variants that fall outside of this range were filtered out.
⍰ Because of the highly repetitive nature of the mitochondrial genome, indel mutations appeared to be more subject to false positives. Therefore, we applied more stringent filtering criteria to indel mutations. For indel mutations to be included, more than half of supporting reads of an indel mutation should have no additional mutations within 10 base pairs.
⍰ Variants that fell in the following regions, which have an extensive level of homopolymers or sequencing error in the reference mtDNA genome (3107N), were explicitly removed:

1. Misalignment due to ACCCCCCCTCCCCC (rCRS 302-315)
2. Misalignment due to GCACACACACACC (rCRS 513-525)
3. Misalignment due to 3107N in rCRS (rCRS 3105-3109)

### Simulating mitochondrial heteroplasmy in maternal line

The mtDNA is present in many copies per cell and is inherited through the maternal germline^78^. In addition, a single cell has different mtDNA variants, and the level of heterogeneity (mitochondrial heteroplasmy) can vary considerably between cells. During the production of primary oocytes, a selected number of mtDNA molecules are transferred into each oocyte. Following the rapid replication of the mtDNA population during oocyte maturation, each oocyte has a variable number of heteroplasmic mitochondrial clones^36^. We attempted to infer the number of clones and levels of mitochondrial heteroplasmy in an oocyte using a simplified random-drift segregation model (**Extended Data Fig. 6b**). This model assumes that (1) a cell has 1,000 copies of mtDNA, and (2) they are partitioned to two daughter cells with an equal amount (500 for each daughter cell), after which (3) all the mtDNA are doubled to reach 1,000 copies of mtDNA in a daughter cells, (4) *f*% of mtDNA in a fertilized egg has the same variant (MT-mtDNA), (5) the variant is neutral, and therefore MT-mtDNA replicates at the same rate as wild type mtDNA (WT-mtDNA). After fifteen generations of cell division, *n*_*samples*_ of cells (the number of clones established in each cadaver) are randomly selected. Two summary statistics were drawn from the *n*_*samples*_ of cells: the proportion of samples harboring MT-mtDNA (*p*), the median heteroplasmic level of MT-mtDNA (*h*). This model simulated 500,000 times for each cadaver, varying the parameter *f* drawn from a uniform distribution with a range of [0.001, 1.000]. We compared the summary statistics (*p, h*) of each simulation to the observed summary statistics, and estimated the posterior distribution of *f* using the neural network regression algorithm of an approximate Bayesian computation^79^.

### Mutational Signature Analysis

We estimated contributions of mutational signatures to an observed mutational spectrum in each sample (i.e., the presumed amount of exposure to corresponding mutational processes) ^41^. This can be achieved by solving the following constrained optimization problem:

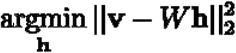

Where 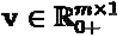, 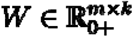, and 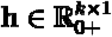 (*m* is the number of mutation types and *k* is the number of mutational signatures). For each sample, given the observed counts of each mutation type **v** from a sample and a pre-trained mutational signature matrix, we calculated exposure **h**. We used an R package (pracma) that internally uses an active-set method to solve the above problem^80^.

The number of mutational signatures is a parameter to be determined for each sample. We started with all known mutational signatures available from the version 3 of the COSMIC mutational signature catalogue and examined the results to sort out the most likely set of signatures (available at https://cancer.sanger.ac.uk/cosmic/signatures). To avoid overfitting, we narrowed them down to a handful of signatures by excluding signatures that do not add much in explaining the mutational spectrum up to a certain fitting error (cosine similarity > ~0.9). Finally, SBS1, SBS5, and SBS18 were used for the analysis of early mutations, and SBS 7a to 7d were additionally considered for the later mutations. For indels, ID1, ID2, ID4, and ID10 for early mutations and ID5, ID8, and ID9 for late mutations were considered.

We found consistent positive correlation between the number of SBS5 mutations () and SBS7a-d mutations ( ) in individuals. We made a linear model estimating (dependent variable) using and age ( ~ + age). Both and age were significant as explanatory variables (both *p* < 2e-16), and the estimate for (*e*) was 0.11. We also estimated the number of SBS18 mutations ( ) using the same variables ( ~ + age). In this analysis, and age was also significant (*p* < 2e-16 and 0.012), and the estimate for (*e’*) was −0.01. Using the above relationships, we calculated the adjusted number of endogenous mutations ( ) as below.

Linear model estimating *n*_*adjENDO*_ by age was more significant than model estimating the number of unadjusted mutations (*n*_*SBS*1_ + *n*_*SBS*5_ + *n*_*SBS*18_) by age (adjusted *R*^2^: 0.46 > 0.17), reflecting effectiveness of the adjustment.

### Copy number variations and structural variations

We identified somatic genomic rearrangements of the whole genome sequenced samples using Delly^81^ with an unmatched blood sample as a control not to miss early mutations existed in matched blood sample. We filtered out the somatic rearrangements in the similar way as our previous report^48^. Briefly, we discarded rearrangement events that were present in the all clonal samples or an in-house panel of normal. Rearrangements with poor mapping quality (Q<25), with an insufficient number of supporting reads (n<5), or those with many discordant reads in matched blood samples were considered to be false positives and were removed. Additionally, we removed short-sized deletions and duplications (<1 Kbp) without soft-clipped reads and unbalanced inversions (<5 Kbp) without sufficient supporting reads (n<5) which are mostly DNA library artifacts. To remove remaining false positive events and to rescue false-negative events located nearby the breakpoints, we visually inspected all the rearrangements passed the filtering process using the IGV^66^.

Segmented copy number profiles were estimated for the whole genome sequenced samples by Sequenza^82^ algorithms using matched blood samples as controls. Copy number changes with length > 10Mbps were included for further analysis. Subclonal copy number changes, of which depth ratios were not properly fitted to the integer values of the absolute copy numbers, were excluded manually.

### Timing estimation of copy number gains

Copy number gains with length > 50Mbps were included in timing analysis. We estimated mutation copy number (*n*_*mut*_) by a previously described formula shown below^67^:

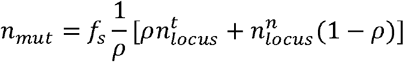

where, *f*_*s*_, *ρ*, 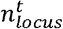, 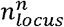 denote variant allele fraction, tumor cellularity, absolute copy numbers in tumor and normal cells, respectively. In our experimental design, the formula will be simplified as follows because *ρ* = 1 (single-cell derived clone):

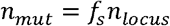

Here, *n*_*locus*_ is given by where and indicates the read depth of the locus of interest in a sample (mean coverage of) and paired bulk blood sample (mean coverage of), respectively.

To assess the temporal relationship between a specific somatic point mutation and chromosomal gains, we compared the mutation copy number ( ) with the allele-specific major copy number ( ) of the chromosomal segment. Mutations with were classified as a pre-amplification event, while the others as post-amplification or minor allele mutations. The probabilities of pre-amplification ( ), post-amplification ( ) were evaluated with the following binomial distributions:

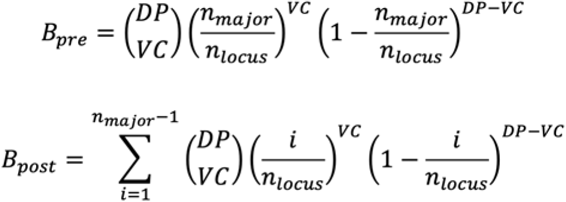

where, and indicate the total read depth and variant read count, respectively. Finally, and are given by:

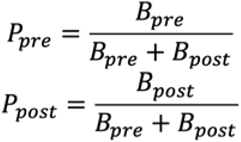

In order to estimate the timing of amplification events, we analyzed the expected number of pre-amplification substitutions ( ) by summation of the values of all substitutions in amplified segments larger than 50Mbp. The 95% confidence interval of was evaluated from z-scores using the sum of as variance. The expected number of post-amplification substitutions ( ) were also calculated in the same way. Then, Age multiplied by the proportion of among clonal mutations ( + were used as chronological timing of copy number gains.

